# P4HA3 is abundantly expressed during early embryonic development, but does not catalyze the formation of 4-hydroxyproline in type I collagen

**DOI:** 10.64898/2025.12.22.695897

**Authors:** Emma Karjalainen, Pekka Rappu, Valerio Izzi, Ilkka Miinalainen, Jyrki Heino, Johanna Myllyharju, Antti M. Salo

## Abstract

4-Hydroxyproline (4Hyp) is crucially important for the collagen triple helix formation and its thermal stability. It is catalyzed by collagen prolyl 4-hydroxylases (C-P4Hs) that are α_2_β_2_ tetramers consisting of two identical catalytic α-subunits (encoded by P4HA1, P4HA2 and P4HA3) and two β-subunits that are protein disulfide isomerases (PDI). The functions of isoenzymes I and II (P4HA1 and P4HA2) are widely studied but very little is currently known about the role of P4HA3. Here we show that *P4ha1* is generally the most abundant isoform in mouse, whereas *P4ha3* expression is tissue and developmental stage specific. *P4ha3* expression is highest in the developing skeletal system and peaks at E12.5 together with the other isoforms. Postnatal *P4ha3* expression is low and mainly present in calvaria, lung, heart and muscle. The *P4ha3* expression virtually disappears at or after 1 week of age in all tissues studied. To evaluate the P4HA3 function in collagen synthesis, we produced cells that lack P4HA3 (*P4ha3−/−*) and cells that have P4HA3 as the only isoform (*P4ha1−/−;P4ha2−/−*) by CRISPR/Cas9. C-P4H activity assays indicated no 4Hyp formation in type I collagen by the *P4ha1−/−;P4ha2−/−* cells, suggesting that the remaining isoform P4HA3 does not participate in type I collagen synthesis. The *P4ha3−/−* cells secreted intact pepsin-resistant collagen that had a similar 4Hyp content and melting temperature as in WT cells, and the secreted collagen assembled correctly into supramolecular collagen fibrils. In conclusion, our results provide new insights into collagen biosynthesis and suggest that prolyl 4-hydroxylation of type I collagen is mediated by P4HA1 and P4HA2 but not by P4HA3.

## Introduction

Collagens are highly abundant proteins in mammals and form a superfamily of 28 collagen types, that have various functions and vital roles, such as maintaining structural integrity of tissues (Gelse et al., 2003; Ishikawa et al., 2025; Kadler et al., 2007; Myllyharju & Kivirikko, 2001; Ricard-Blum, 2011). Collagen molecules are formed by three α chains that have a repeating -Gly-X-Y-amino acid sequence and are folded into a triple helix. The X and Y positions can accommodate any amino acid, but the X position is frequently proline and Y is 4-hydroxyproline (4Hyp). 4Hyp is a characteristic feature of collagen molecules as the majority of Y position prolines are hydroxylated. 4Hyp is essential for folding of the collagen triple helix and its thermal stability, and the melting temperature of the triple helix reflects the amount of 4Hyp (Bächinger & Boudko, 2025; Gelse et al., 2003; Myllyharju & Kivirikko, 2004; Salo & Myllyharju, 2021; Shoulders & Raines, 2009).

Formation of 4Hyp in procollagen polypeptides is catalyzed by prolyl 4-hydroxylases (C-P4Hs, EC 1.14.11.2) in the lumen of the endoplasmic reticulum (ER). C-P4Hs are 2-oxoglutarate (2OG) - dependent dioxygenases, which require 2OG, Fe^2+^, ascorbate and oxygen in the reaction mechanism (Myllyharju, 2003a, 2015; Salo & Myllyharju, 2021). C-P4Hs are α_2_β_2_ tetramers with three isoforms of the catalytic α-subunit (encoded by *P4HA1, P4HA2* and *P4HA3*) and a β-subunit is identical to protein disulfide isomerase (PDI, encoded by *P4HB*) that form the C-P4H-I, C-P4H-II and C-P4H-III isoenzymes, respectively (Annunen et al., 1997; Helaakoski et al., 1995; Kukkola et al., 2004; Myllyharju & Kivirikko, 2004). PDI employs its active site cysteines to bind with an α-subunit (Murthy et al., 2018, 2022a) and keeps the otherwise insoluble α-subunits soluble and provides a KDEL ER retention sequence for the C-P4H tetramer (Helaakoski et al., 1995; Pihlajaniemi et al., 1987; Vuori et al., 1992; Wilson et al., 1998). The C-P4H α-subunit has an N-terminal dimerization domain followed by a peptide substrate-binding (PSB) domain, and a C-terminal catalytic (CAT) domain (Anantharajan et al., 2013; Koski et al., 2017; Myllyharju & Kivirikko, 1999). A small-angle X-ray scattering data suggests that the C-P4H tetramer has an elongated symmetric βααβ assembly (Koski et al., 2017). The two α-subunits bind to each other from their N-terminal coiled coil domains, while the PSB domains stretch into opposite directions and the CAT domain and PDI/β-subunit interact closely at the ends of the tetramer (Koski et al., 2017; Murthy et al., 2022b).

C-P4H-I is the main isoform (Holster et al., 2007; Salo et al., 2024; Zou et al., 2017) and is essential for viability as *P4ha1*^−/−^ mice die shortly after embryonic day 10.5 due to ruptures in the basement membrane caused by abnormal assembly of type IV collagen (Holster et al., 2007). *P4ha2*^−/−^ mice are viable and do not have any obvious phenotypic abnormalities, but interestingly, when combined with *P4ha1* haploinsufficiency (*P4ha1*^*+/-*^; *P4ha2*^−/−^ mice) a significant reduction in total C-P4H activity is observed with consequent defects in collagen quality, properties of cartilaginous ECM, growth plate survival and proper formation of bone matrix (Aro et al., 2015; Tolonen et al., 2022). Currently, not much is known about the C-P4H-III. P4HA3 was identified and characterized in 2003 (Kukkola et al., 2003; Van Den Diepstraten et al., 2003) and P4HA3 was shown to be expressed in various human tissues, but in lower levels when compared to the other isoforms (Kukkola et al., 20023), strongest expression being found in the fibrous cap of atherosclerotic plaques (Van Den Diepstraten et al., 2003). P4HA3 has been suggested to form an active C-P4H with PDI with similar catalytic properties to C-P4H-I and II (Kukkola et al., 2003). Several studies have shown that expression of P4HA3 is upregulated in various cancers and it is involved in cancer progression. For example, P4HA3 expression is upregulated in melanoma, and it promotes tumor proliferation and invasion (Atkinson et al., 2019). In gastric cancer, P4HA3 is significantly upregulated when compared to normal stomach tissue and it is suggested to enhance gastric cancer cell motility and invasiveness, high P4HA3 expression hence correlating with poor overall survival (Song et al., 2018). Renal cell carcinoma (RCC) patients have been shown to have a higher P4HA3 expression, and silencing of P4HA3 inhibits RCC cell proliferation, migration, and invasion (Shi et al., 2021). P4HA3 is upregulated also in head and neck squamous cell carcinoma and promoted cell proliferation, migration, and invasion by inducing epithelial-mesenchymal transformation (Wang et al., 2020). Recently, collagen sequence specificity for C-P4H-I and C-P4H-II was discovered. C-P4H-I preferentially hydroxylates XPG triplets with positive and polar uncharged amino acids in the X-position (Salo et al., 2024), whereas the biological function of C-P4H-II is to hydroxylate collagen EPG and DPG sites (Salo et al., 2024; Wilhelm et al., 2023) that C-P4H-I hydroxylates poorly or not at all (Salo et al., 2024). The data also shows that this selectivity arises from differences in the active sites of the two C-P4H isoenzymes (Salo et al. 2024a). Here, our aim was to study what is the role of C-P4H-III in the hydroxylation of the most abundant collagen, the type I collagen.

## Materials and methods

### Cell culture and generation of knockout cell lines

Cell lines used in these experiments were Large T immortalized WT and *P4ha1*^−/−^ mouse embryonic fibroblasts (MEFs) from 10.5 dpc (Holster et al., 2007), and MC3T3 preosteoblasts. MEFs were cultured in Dulbecco’s modified Eagle Medium (DMEM) + GlutaMAX™-I (31966-021, Gibco) supplemented with 10% fetal bovine serum (FBS, Sigma-Aldrich) and 100 U/ml penicillin with 0.1 mg/ml streptomycin (Sigma-Aldrich). MC3T3 cells were cultured in Minimum Essential Medium (MEM) Eagle Alpha, w Stable L-Glutamine w Deoxyribonucleosides & Ribonucleosides (ECM0850L, Euroclone) supplemented with 10 % FBS (Sigma-Aldrich) and 100 U/ml penicillin with 0.1 mg/ml streptomycin (Sigma-Aldrich). Mouse primary brain microvascular endothelial cells were cultured in endothelial cell basal medium (210-485, Cell Biologics) with appropriate supplements (M1168-Kit, Cell Biologics). 10 cm culture plates were precoated with gelatin-based coating solution (6950, Cell Biologics) before seeding the endothelial cells. For collagen extraction analyses, FBS was replaced with a protein-free replacement, Panexin CD (P04-930500, PAN-Biotech) to avoid any trace of bovine collagen. All cells were incubated at 37°C in a humidified atmosphere of 5 % CO2 and 95 % air. CRISPR construct design was done with the CRISPOR.org web tool (Concordet & Haeussler, 2018). A pSpCas9(BB)-2A-GFP plasmid was a gift from Feng Zhang (Addgene plasmid # 48138; http://n2t.net/addgene:48138; (Ran et al., 2013)) and was used as a backbone for the CRISPR/Cas9 constructs and the guide-RNAs were cloned between the U6 promoter and the scaffold of gRNA in PX458 plasmid as follows. A non-targeting, scrambled sequence 5’-GCACTACCAGAGCTAACTCA-3’ (OriGene) was used as a negative control. First, sense and antisense oligos (100 µM, Sigma-Aldrich) (Table S1) were annealed using 1 M Tris-HCl (pH 8). The annealing mixtures were heated in boiling water in 500 ml Erlenmeyer flask and allowed to cool down overnight. Next, the PX458 plasmid was digested with BpiI/BbsI (ER1011, Thermo Fisher Scientific) restriction enzyme that recognizes GAAGAC(2/6)^ sites resulting in a 9000-bp long linear fragment. The digestion mix was incubated for 1 hour at 37°C and then separated on a 0.8% agarose gel in 40 mM Tris, 20 mM acetic acid, 1 mM EDTA-buffer (TAE). The gel was visualized using ChemiDoc XRS+ gel imager (Bio-Rad). The digested plasmid gel piece was cut on UV-board and purified using Illustra™ GFX PCR DNA and Gel Purification Kit (28-9034-70, GE Healthcare) as per manufacturer’s instructions. The annealed oligo mixes were diluted to 1:200 with H_2_O and then inserted into the digested plasmid (100 ng) using Quick T4 ligase (New England Biolabs). The correct insertion of gRNA into the CRISPR plasmid was confirmed by sequencing, using five pmol of U6 sequencing primer GACTATCATATGCTTACCGT-3’ and 200-250 ng of each purified plasmid. Sequencing was performed by using ABI3500xL Genetic Analyzer in the Biocenter Oulu Core Facility. Sequence-verified colonies were chosen and inoculated into a maxi-prep culture and plasmid DNA was purified using QIAprep midi Kit (12143, Qiagen) as in manufacturer’s instructions. After confirming that the CRISPR/Cas9 constructs were correct, the plasmids were transfected to WT and *P4ha1*^−/−^ MEFs and WT MC3T3s. Lipofectamine 3000 (L3000-008, Invitrogen) was used to deliver gRNA plasmids (10ng each) according to the manufacturer’s instructions. PX458-P4ha3-ex2 plasmid was transfected to WT MEFs and MC3T3s to create a *P4ha3*^−/−^ clones, and PX458-P4ha2-ex3 plasmid was transfected to *P4ha1*^−/−^ cells to create a *P4ha1*^−/−^*;P4ha2*^−/−^ MEFs. WT and *P4ha1*^−/−^ MEFs and WT MC3T3s were also transfected with scrambled plasmid to create control single cell clones. Fluorescence-activated cell sorting (FACS) analysis was used to isolate single cell clones of the transfected cell lines. Cell sorting was performed with BD FACSAria IIIu sorter at the Translational Cell Biology Core Facility of the Kontinkangas campus, University of Oulu. Before sorting, cells were detached from the plates using trypsin-EDTA (MS0158100U, Biowest), centrifuged, and resuspended in fresh PBS supplemented with 1% FBS to obtain a concentration of 3×106 cells/ml. The cell suspensions were filtered using 35-µm cell strainer and 7-aminoactinomycin D staining solution was added to discriminate live cells from dead cells. Single cells were sorted onto individual wells of a 96-well plate. Single cell clones were expanded in 24-well, 6-well, and 10-cm plates until cell numbers were sufficient for cell banking, genomic DNA extraction, western blot and qPCR analyses, and further experiments.

### Cell line validation

Genomic DNA was extracted using QuickExtract DNA solution 1.0 (QE0905T, Lucigen) from 24-well plate cell cultures. Cells were washed with PBS and 50 μl of QE solution was added to the wells. Cells were scraped and subsequently incubated for 15 min at 65°C, 15 min at 68°C and 10 min at 98°C. The Genomic DNA was amplified using Phusion High-Fidelity DNA polymerase (Thermo Fisher Scientific) and gene-specific primers (Table S2). Half of the PCR products were digested with specific restriction enzymes (New England Biolabs). For P4ha2-ex2, the used restriction enzyme was BsoMI (R0586S) and for P4ha3-ex2 BsrI (R0527S). Digested and undigested PCR products were separated on a 2 % agarose gel and imaged using ChemiDoc™ XRS+ -gel imager (Bio-Rad). Putative positive samples were cut from the gel and purified using GFX PCR DNA and Gel Purification Kit (GE Healthcare). The purified PCR products were sequenced using 5 pmol of primer and approximately 3-20 ng DNA using an ABI3500xL Genetic Analyzer in the Biocenter Oulu Sequencing Core Facility. Single cell clones were cultured until confluency and then washed with PBS and lysed using a lysis buffer (0.1 M NaCl, 0.1 M glycine, 10 mM Tris, 0.1% Triton X-100, 1x phosphatase inhibitor (P5726-1ML, Sigma) and 1x EDTA-free protease inhibitor (04693132001, Roche) by 3x freezing/thawing cycles and samples were frozen at −70°C. Lysed cells were centrifuged at 10 000 x g for 15 min at 4°C to separate insoluble and soluble protein fractions. Protein concentration was determined from soluble fraction using Direct Detect Spectrophotometer (Millipore Corporation). Typically, 30-40 µg (70 µg in case of P4HA3 analysis) of the soluble fraction protein lysates were resolved under reducing condition in 8% sodium dodecyl sulfate-polyacrylamide (SDS-PAGE) gel or under nonreducing and nondenaturing conditions on 8% native PAGE gel without SDS and reducing agent. The gel was electroblotted onto polyvinylidene difluoride (PVDF) membrane (Immobilon P, Millipore). Proteins were detected with either Pierce™ ECL Western Blotting Substrate (32106, Thermo Fisher Scientific), Immobilon chemiluminescent HRPsubstrate (WBKLS0100, Millipore) or Clarity Max Western ECL substrate (1705062, Bio-Rad) and imaged using Molecular Imager ChemiDoc™ XRS+ (Bio-Rad). Primary antibodies were rabbit polyclonal anti-P4HA1 (1:500, A3999, ABclonal), rabbit polyclonal anti-P4HA2 (1:500, 13759-1-AP, Proteintech), rabbit polyclonal anti-P4HA3 (1:500, HPA007897, Sigma), and mouse monoclonal anti-β-actin (1:20000, NB600-51, Novus). Horseradish peroxidase-conjugated (HRP) anti-mouse or anti-rabbit were used as secondary antibodies (1:10000, DAKO).

### Expression analysis

Total RNA was extracted from single cell clones using E.Z.N.A^®^ Total RNA Kit I (6834-01, Omega Bio-Tek) according to manufacturer’s protocol. RNA was treated with RNase free DNase (E1091-02, Omega Bio-Tek). Total RNA concentrations were measured by Nanodrop™ spectrophotometer (ThermoFisher). The iScript cDNA synthesis kit (1708890 or 1708891, Bio-Rad) was used for reverse transcription according to the manufacturer’s instructions. WT C57BL6/N male mice were euthanized at the following time points: newborn (P0), day 2 (P2), day 4 (P4), 1, 2, and 6⍰weeks and various tissues were harvested. In addition, fetuses from different development days were collected. MEFs, MC3T3 cells and mouse brain microvascular endothelial cells were harvested at 70-80% confluency. Total RNA was isolated using either E.Z.N.A total RNA kit I (Omega Bio-tek) or Trizol (Thermo Fisher Scientific). The iScript cDNA synthesis kit (Bio-Rad) was used for reverse transcription according to the manufacturer’s instructions. A QX200™ Droplet Digital PCR system (Bio-Rad) was used for analysis of expression of the *P4ha1, P4ha2* and *P4ha3* genes. For each reaction, 2x ddPCR Supermix for probes (Bio-Rad) was mixed with 250 nM TaqMan Gene Expression Assay (FAM) components (Thermo Fisher Scientific) and the prepared cDNA template (33 ng) to a final volume of 22 µl. The TaqMan Gene Expression Assays (FAM) used were P4ha1 FAM (Mm00803137_m1), P4ha2 FAM (Mm00477940_m1) and P4ha3 FAM (Mm00622868_m1). 20 µl of the samples were loaded onto an eight-channel cartridge (Bio-Rad) along with 70 µl of droplet generation oil for probes (Bio-Rad). Following emulsion generation on the QX200™ Droplet Generator (Bio-Rad), samples were transferred to a 96-well PCR plate, heat-sealed with foil (1814040, Bio-Rad) by using PX1™ PCR plate Sealer (Bio-Rad), and amplified in T100™ Thermal Cycler (Bio-Rad). Thermal cycling conditions were 95°C for 10 min, followed by 40 cycles of 94°C for 30 s, 60°C for 1 min, and 98°C for 10 min with a ramp rate of 2°C/s. Samples were kept overnight at 4°C. Next day, droplets were analyzed with QX 200™ Droplet Reader (Bio-Rad). Results were reported as copies per µl of reaction, determined by the QuantaSoft™ software (Bio-Rad). Absolute expression levels were shown as copies per µg of input RNA and relative expression levels of *P4ha1, P4ha2* and *P4ha3* were calculated from the results and as percentage (%) ± SD. For quantitative PCR, cells were seeded on 6-well plates in a density of 2.5×10^5^ cells/well and total RNA was extracted by using E.Z.N.A^®^ Total RNA Kit I (R6834-01, Omega Bio-tek) according to the manufacturer’s protocol and treated with RNase free DNase (E1091-02, Omega Bio-Tek). Total RNA concentrations were measured by Nanodrop™ spectrophotometer (Thermo Fisher Scientific). The iScript cDNA synthesis kit (1708890 or 1708891, Bio-Rad) was used for reverse transcription according to the manufacturer’s instructions. Quantitative real-time PCR (qPCR) analysis was performed using iTaq Universal SYBR Green Supermix (1725124, Bio-Rad) according to the manufacturer’s instructions with a CFX96 thermal cycler. Expression levels were normalized to the geometrical mean of TATA-box binding protein (Tbp), glyceraldehyde-3-phosphate dehydrogenase (*Gapdh*) and β-actin (*Actb*) mRNA levels using the comparative Ct (2^-ΔCt^) method. The primers used for qPCR analysis are listed in Table S3.

### Open data (OA) analysis

In situ hybridization (ISH) images were retrieved from the EMAGE gene expression database (http://www.emouseatlas.org/emage/) (Richardson et al., 2014). *P4ha3* is shown as pseudo-wholemount for enhanced clarity and expression regions and structures were downloaded and summarized. Single cell RNAseq (scRNAseq) data of mouse embryonic development from gastrula to birth (Qiu et al., 2024) were downloaded from the CZ CELLxGENE Discover platform and reanalyzed as reported by the authors (https://github.com/ChengxiangQiu/JAX_code) in R/Python.

### Collagen extraction

Cells were seeded on two 15 cm dishes with a density of 1.5×10^6^ cells/plate. After two days, cells were washed twice with PBS and serum-free culture medium containing 10% Panexin CD (P04-930500, Pan Biotech) was added. Following day, medium was exchanged and 150 μg/ml of L-ascorbic acid phosphate magnesium salt n-hydrate (Wako) was added on two consecutive days and medium was collected on the third day. The medium collection procedure was repeated and on the second time also the cells were collected. Collagen extraction from the medium and cell samples was done with and without pepsin digestion as previously described (Miller & Rhodes, 1982) with slight modifications. Triple-helical collagen is resistant to pepsin and thus pepsin digestion was used to analyze the presence of folded triple-helical collagen molecules. The two collected medium samples were combined and centrifuged at 3500 x g for 30 min at +4°C to remove any dead cells and the supernatant was transferred to a new tube. Collagen precipitation was done by adding 176 mg/ml of ammonium sulphate to the medium and incubated overnight with mixing at +4°C. Samples were centrifuged at 3500 x g for 45 min at +4°C, supernatant was discarded, and the pellet was dissolved in 0.5 M acetic acid with 0.1 mg/ml pepsin (10108057001, Roche Diagnostics) and incubated overnight with mixing at +4°C. The following day, samples were centrifuged at 20000 x g for 30 min at +4°C and supernatant was collected and precipitated by adding 4 M NaCl to a final 0.7 M concentration. Samples were incubated overnight with mixing at +4°C, centrifuged at 20000 x g for 30 min at +4°C, supernatant was discarded, and the collagen pellet was dissolved in 0.1 M acetic acid. Aliquots (10µl) of the collagen samples were analyzed with SDS-PAGE. The pH of the collagen samples was neutralized with 1.5 M Tris-HCl (pH 8.8), and samples were analyzed by 6% SDS-PAGE under reducing conditions. The proteins were stained with Coomassie Brilliant Blue R-250 followed by destaining with 10% % EtOH, 10 % CH_3_COOH in H_2_O, and imaged with a Gel Doc™ XRS+ molecular imager (Bio-Rad).

### Amino acid analysis

Aliquots (10 µl) of the collagen samples were resolved under reducing condition in 8% SDS-PAGE and transferred onto a ProBlott™ membrane (400994, Applied Biosystems) as per manufacturer’s instructions. Protein bands from the membrane were cut into small pieces, washed sequentially with 0.2% triethanolamine in 50% methanol, 100% methanol and dH2O, and then transferred to prepyrolyzed hydrolysis tubes (6×60 mm, Duram). The tubes were inserted into a 50-ml hydrolysis vial containing 1 ml of 5.7 N hydrochloric acid solution (Sigma-Aldrich) and purged with nitrogen. The hydrolysis was performed at +150°C for 75 min. After hydrolysis, the sample membrane was wet with 15 µl of methanol and the hydrolyzed amino acids were extracted with 35 µl of 0.1 M HCl at +4°C overnight. Each sample was transferred to an Eppendorf tube and dried completely with a Speed-Vac overnight. The hydrolysate from each tube was dissolved in 50 μl of 20 mm HCl and then filtered 61 with an Ultrafree-MC (0.45 μm, Millipore). Amino acid derivatization was performed with a Waters AccQ·Tag chemistry package according to the manufacturer’s instructions. The derivatized amino acids were analyzed with a Shimadzu Prominence HPLC with its fluorescence detector set at 250/395 nm at the Biocenter Oulu Proteomics and Protein Analysis Core Facility.

### Circular dichroism spectroscopy

To evaluate whether P4HA3 deficiency affects the melting temperature of the collagen molecules, aliquots (10 µl) of the medium collagen samples were analyzed by circular dichroism (CD) spectroscopy using a Chirascan CD spectrometer (Applied Photophysics). The collagen samples were diluted with 0.1 M acetic acid to a concentration of 0.1 mg/ml. The concentration of each sample was verified with absorbance at 205 nm. CD data were collected between 280 and 190 nm at +20°C using a 0.1-cm path-length quartz cuvette. CD measurements were acquired every 1 nm with 0.5 s as an integration time and repeated three times with baseline correction. Data were processed using Chirascan Pro-Data Viewer (Applied Photophysics). The direct CD measurements (θ, mdeg) were converted into mean residue molar ellipticity ([θ]MR) by Pro-Data Viewer. Thermal denaturation of the protein samples was monitored by measuring the CD spectra in the same setup with a temperature range from +20°C to +96ºC at a rate of 1°C/min using a Peltier 63 Temperature Control TC125 (Quantum Northwest). The CD data were recorded at every 2°C. The melting temperature (Tm) of the triple helix was calculated with GraphPad Prism 10.1.2 (GraphPad Software).

### Mass Spectrometric analysis

The relative amount of proline and 4-hydroxyproline from collagen (Table S4) was analyzed by mass spectrometric method. First, concentrations of purified medium collagens were measured with Direct Detect infrared spectrophotometer (Millipore Corporation) and 20 µg of purified medium collagen samples were vacuum-dried with SpeedVac. The proteins in dried gel pieces were alkylated and digested as described (Shevchenko et al., 2006) except that trypsin-LysC mix (Promega) was used instead of trypsin, and the eluted peptides were redissolved in 0.1% formic acid instead of 0.1% trifluoroacetic acid. The peptides were loaded on a nanoflow HPLC system (Easy-nLC1000, Thermo Fisher Scientific) coupled to the Q Exactive HF mass spectrometer (Thermo Fisher Scientific) equipped with a nano-electrospray ionization source. The peptides were first loaded on a trapping column and subsequently separated in-line on a 15 cm C18 column (75 µm × 15 cm, ReproSil-Pur 3 µm 200 Å C18-AQ, Dr. Maisch HPLC GmbH, Ammerbuch-Entringen, Germany). The mobile phase consisted of water/acetonitrile (98:2 (v/v)) with 0.2% formic acid (solvent A) and acetonitrile/water (95:5 (v/v)) with 0.2% formic acid (solvent B). The peptides were separated with a 10-min gradient from 4 to 30% of solvent B. Before the end of the run, the percentage of solvent B was raised to 100% in 2 min and kept there for 8 min. Full MS scan over the mass-to-charge (m/z) range of 300-2000 with a resolution of 120,000, followed by data dependent acquisition with an isolation window of 2.0 m/z and a dynamic exclusion time of 10 s was performed. The top 6 ions were fragmented by higher energy collisional dissociation with a normalized collision energy of 27% and scanned over the m/z range of 200-2000 with a resolution of 15,000. After the MS2 scan for each of the top 6 ions had been obtained, a new full mass spectrum scan was acquired and the process repeated until the end of the run. Tandem mass spectra were searched against a mouse reference proteome from UniProt (database version 2021_4) with Proteome Discoverer software, version 2.4 (Thermo Fisher Scientific) using MS Amanda 2.0 search engine (Dorfer et al., 2014) allowing for 7 ppm precursor mass tolerance and 0.02 Da fragment mass tolerance. Carbamidomethyl (C) as a fixed modification, and oxidation (M, K, P) as dynamic modifications were included. Maximum of two missed cleavages were allowed. Decoy database search using reversed sequences was used to assess the false discovery rate. Only the peptide-spectrum matches with false discovery rate < 0.05 were accepted. The list of spectra matching to collagen-derived peptides was used to calculate relative proline hydroxylation of all possible XPG-triplets. Relative total hydroxylation for each PG sequences was obtained by dividing the number of MS/MS spectra matching to any peptide containing only hydroxylated PG sequence(s) with the number of MS/MS spectra matching to any peptide containing at least one hydroxylated or non-hydroxylated PG sequence.

### Collagen prolyl 4-hydroxylase activity assay

C-P4H activity/µg of soluble protein of the cells was assayed by a method based on the formation of 4-hydroxy[^14^C]proline in a [^14^C]proline-labelled protein substrate consisting of non-hydroxylated chick type I protocollagen chains (Kivirikko & Myllylä, 1982). The cell protein samples were prepared as described for cell line validation. Protein concentrations were measured using Direct Detect infrared spectrophotometer (Millipore Corporation). Briefly, cell lysates were incubated with the [^14^C]proline-labelled substrate, which is denatured by boiling, and co-factor mix (0.5 M Tris-HCl, pH7.8, 20 mg/ml BSA, 44500 U/mg protein catalase, 0.01 M DTT, 1.46 mg/ml α-ketoglutarate, 3.52 mg/ml ascorbate, 0.278 mg/ml FeSO_4_) at 37°C for 30 min. The reaction was stopped by adding 6 M HCl and samples were hydrolyzed (Kivirikko & Myllylä, 1982). The formed radioactively labelled 4Hyp was then measured in a scintillation counter (Liquid Scintillation Analyzer, Tri-Carb 2900TR, PerkinElmer) after transforming it to pyrrol using chloramine-T as an oxidizing agent (Berg, 1982). The results were reported as a mean DPM/μg ± SD. *In vitro* production of human procollagen α(I) was done as previously (Salo et al., 2024). Briefly, TNT T7/SP6 coupled reticulocyte lysate system (Promega) was used to produce substrate in tubes that contained dried L-[2,3,4,5-3H]-Proline. The unbound proline was dialyzed out and radiolabeled procollagen was used as a substrate in the reaction with recombinant C-P4H-I as a positive control and WT, *P4ha3*^−/−^, *P4ha1*^−/−^*;P4ha2*^−/−^ lysates as a source of enzyme to measure 4Hyp as described above.

### Transmission electron microscopy

Cells were seeded on UpCell™ surface plates (174899, Thermo Fisher Scientific) and grown for 5–7 days in a culture medium containing 150 µg/ml L-ascorbic acid phosphate magnesium salt n-hydrate (Wako). The cells were detached from the plates according to the manufacturer’s instructions and fixed using 1% glutaraldehyde, 4 % paraformaldehyde in 0.1 M phosphate buffer (pH 7.2) for transmission electron microscopy (TEM). The samples were post-fixed in 1% osmium tetroxide, dehydrated in acetone and embedded in Epon LX 112 (Ladd Research Industries, Vermont, USA). Thin sections (70 nm) were cut with Leica Ultracut UCT ultramicrotome and stained in uranyl acetate and lead citrate. Samples were examined in a Tecnai G2 Spirit 120 kV TEM (FEI Europe, Eindhoven, Netherlands) operated at 100 kV and images were captured by Quemesa CCD camera (Olympus Soft Imaging Solutions GmbH, Münster, Germany) using Radius software (EMSIS GmbH, Münster, Germany). Analysis of images was done using ImageJ software (Fiji). The collagen fibril diameter was measured from WT, *P4ha1*^−/−^ and *P4ha3*^−/−^ MEFs, and WT and *P4ha3*^−/−^ MC3T3 cells, and the mean diameter per each genotype was calculated. The results were recorded as a mean ± SD.

### Statistical analyses

GraphPad Prism (Version 10.1.2, GraphPad Software, Inc.) was used for all statistical analyses. The data in the current study are presented as mean ± SD. A two-tailed student’s t-test was used to determine statistical significance between two groups and ordinary one-way ANOVA was used with more than two groups. P-value <0.05 was considered statistically significant. Three standard deviations from the mean were considered an outlier.

## Results

### Expression of *P4ha* isoforms during mouse development

To investigate the role of P4HA3 we first studied the absolute and relative mRNA expression levels of the three *P4ha* isoforms, the emphasis being on the analysis of *P4ha3* expression. We used ddPCR to measure the absolute mRNA levels of *P4ha1, P4ha2* and *P4ha3* during mouse embryonic development and in postnatal mouse tissues. First, the absolute expression levels were measured from the whole mouse embryos at various time points (Fig. 1). *P4ha1* was found to be the most abundant isoform in all time points contributing more than 60% of the *P4ha* transcripts and *P4ha2* was generally the second most abundant isoform (Fig. 1). However, the *P4ha3* expression was higher than the *P4ha2* between the embryonic days 11.5-13.5 dpc. In the time-window that was used in the analysis (8.5-18.5 dpc), the highest expression of all isoforms was detected at 12.5 dpc (Fig. 1A), the relative expression of *P4ha1* being 61%. *P4ha2* 16% and *P4ha3* contributed 23% to the total amount of *P4ha* mRNA (Fig. 1B). Previously we have shown the *P4ha3* expression in the developing mice in bone and growth plate (Tolonen et al., 2022). Now, we performed a more complete expression analysis of various mouse tissues at different post-natal time points. Again, we used ddPCR to analyze the absolute transcript levels (Fig.2). Generally, the absolute expression values of all three isoforms were at highest in the early time points, typically peaking at P2 and P4, and then decreasing substantially by the 2-week time point (Fig. 2). However, in some tissues, e.g. the kidney and liver, the expression levels were relatively similar in all time-points from P0 to 6wk old. In some tissues, e.g. the lung and heart, expression of *P4ha1* and *P4ha2* started to increase again at the 6-week time point. As in the embryos (Fig. 1), *P4ha1* was found to be the most abundant isoform in the post-natal tissues with only a few exceptions (Fig. 2). In 1-week lung tissues, *P4ha3* expression surpassed the *P4ha1* as well as the *P4ha2* expression. Similarly, we have observed previously that in P0 growth plates, *P4ha3* expression surpassed the *P4ha1* expression, while *P4ha2* was the most abundant isoform in the distal epiphysis of the 6-week femur (Tolonen et al., 2022). *P4ha3* typically had its highest expression levels during the early post-natal days with a notable decrease at the age between 2 to 6 weeks. Similarly to previous study (Tolonen et al., 2022), high absolute *P4ha3* expression level was detected in calvaria, up to 400 copies/ng of input RNA in the early post-natal time points. In calvaria, *P4ha3* also had a relatively high expression among the isoforms until the 1–2-week time points. The second highest absolute *P4ha3* expression levels, about 200 copies/ng of input RNA, were detected in the lung (as the most abundant isoform at week 1), heart (at P2) and calf muscle (at P2 and P4). In all other tissues and time points the absolute expression levels of the *P4ha3* transcript were markedly lower or even undetectable. In addition, we plotted the results as relative expression values (Fig. S1) to visualize the contribution of each isoform. The expression of *P4ha* isoforms followed similar pattern in all investigated tissues. *P4ha1* was relatively the most abundant isoform. *P4ha2* or *P4ha3* are relatively high compared to *P4ha1* during the early development time points and then decreases at age. Interestingly the expression of *P4ha3* in lung is relatively high in early time points, and at 1wk time point *P4ha3* became the expressed isoform (50%). At 6-week time point, the relative expression of *P4ha3* has decreased drastically, whereas *P4ha1* remained the main isoform and *P4ha2* the second highest isoform. We also analyzed the total amount of *P4ha* mRNA, i.e. all three isoform transcripts combined, in the postnatal tissues (Table S5). Overall, the highest absolute total *P4ha* expression values were observed in the bones, growth plate, lung, eye, kidney, brain, heart and calf muscle. The highest total *P4ha* transcript values were typically detected at the early developmental time points and then decreased by the 2–6-week time point. Markedly lower amounts were observed in the skin, liver, spleen and intestine (Table S5). In the tendon samples extracted from different tissues of 6-week-old mice, the total *P4ha* mRNA amount was also high in the tail tendon (Table S5), resulting almost exclusively from *P4ha1* expression (Fig. 2, Table S5), whereas in the flexor digitorum longus (FDL) and Achilles tendons the *P4ha* isoform expressions were low. *P4ha3* expression was extremely low or non-existent in all tendon samples (Fig. 2). To validate the expression patterns of *P4ha3*, we compared our findings with comprehensive datasets profiling mouse embryonic development, including in situ hybridization (ISH) and single-cell RNA sequencing (scRNA-seq). ISH data confirmed the spatial expression pattern of *P4ha3* across multiple embryonic tissues (Fig. 3A). *P4ha3* was robustly expressed in distinct anatomical structures, spanning a broad array of physiological systems, with particularly high expression within the developing skeletal system (Fig. 3A, Fig S2). Consistent with these findings, analysis of a scRNA-seq atlas comprising over 11 million cells from gastrulation to birth (Fig. 3B, Fig. 3C, Fig. S3 and Fig. S4) confirmed prominent *P4ha3* expression in neural and neuroectodermal precursors, mesodermal populations, epithelial and endothelial cells, and hematopoietic lineages. Within the mesoderm, expression was especially enriched in facial mesenchyme, lateral plate and intermediate mesoderm, sclerotome, and limb progenitor populations, closely mirroring the ISH results.

**Figure 1.**
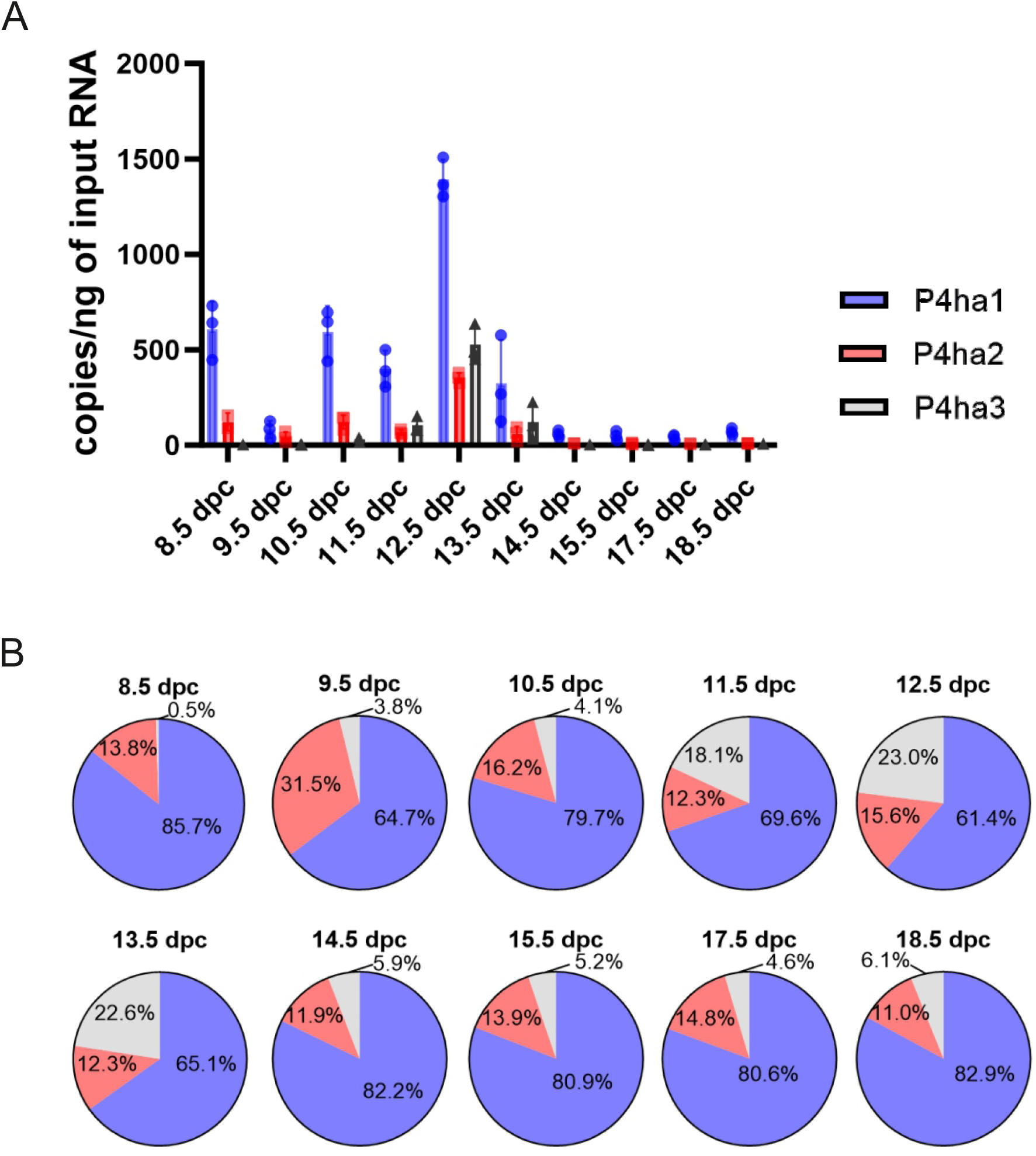
Droplet digital PCR analysis of *P4ha1, P4ha2* and *P4ha3* transcripts in WT C57BL/6N mouse embryos at various time points between 8.5 dpc and 18.5 dpc shown as (A) copies/ng of input RNA and (B) relative abundance, n=3 biological replicates per time point.

**Figure 2.**
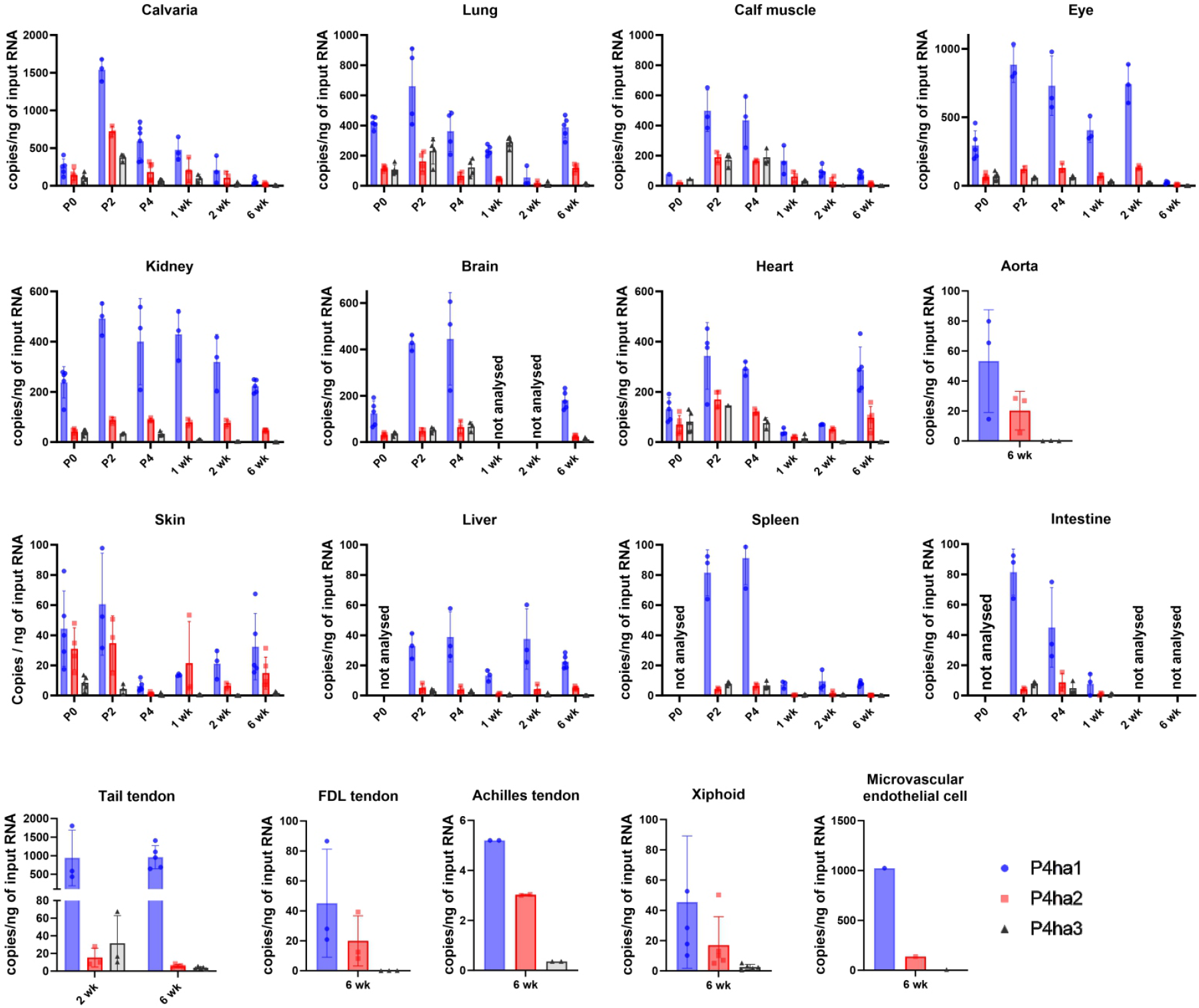
Droplet digital PCR analysis of *P4ha1, P4ha2* and *P4ha3* transcripts in postnatal tissues. The data are shown as absolute expression values, copies/ng of input RNA. Tissues are from newborn (P0) to 6-week-old WT C57BL/6N mice, n=3–5 biological replicates per time point.

**Figure 3.**
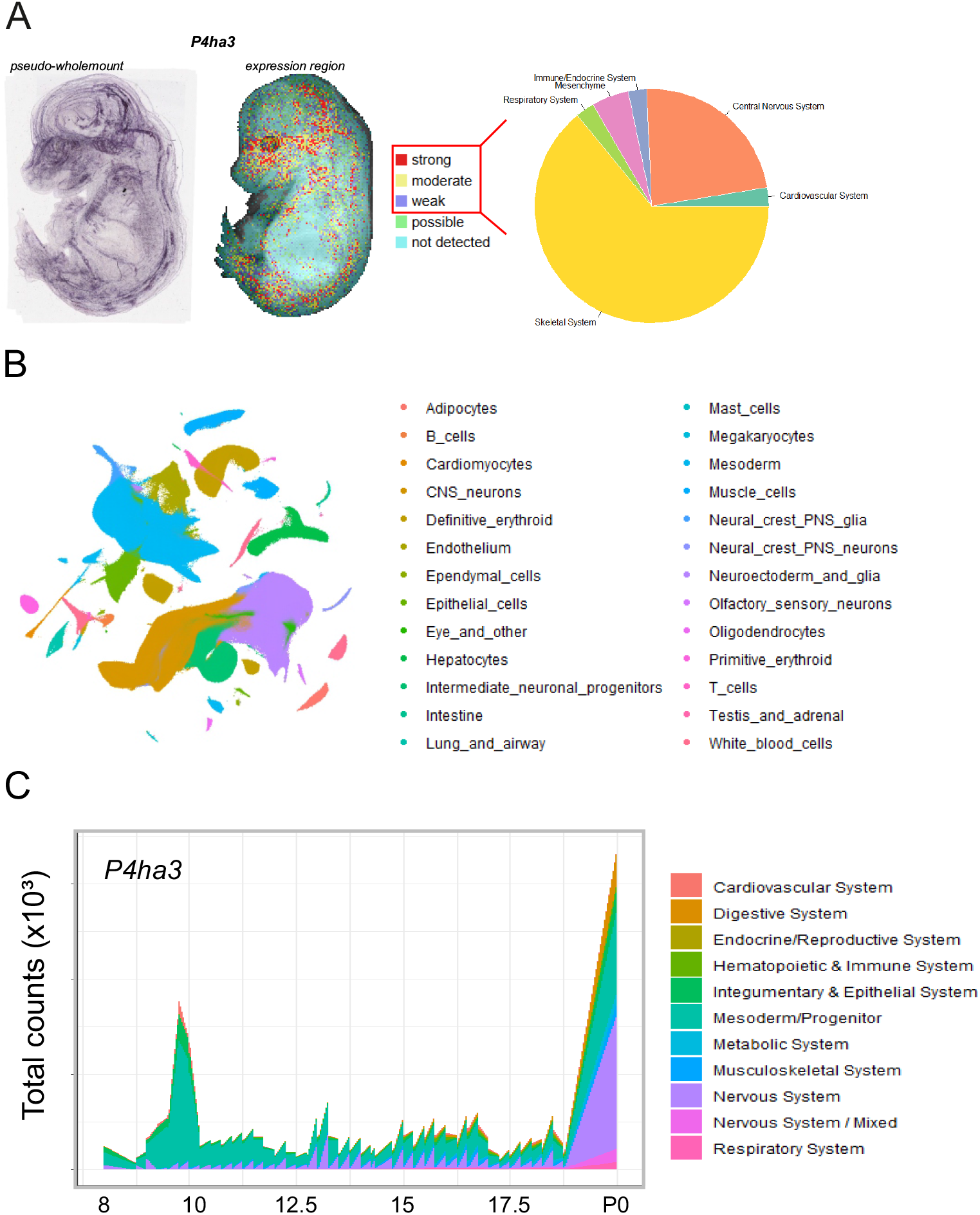
*P4ha3* expression in OA data. Analysis of open-access (OA) data from the in situ hybridization (ISH) collection at (A) EMAGE (Richardson et al., 2014) and (B) scRNA-seq from (Qiu et al., 2024) show strong mesenchymal expression of *P4ha3*, with particularly high abundance in the skeletal system and (C) changing spatiotemporal patterns across development phase.

### Generation and validation of C-P4H deficient cell lines

To study the role of P4HA3 in collagen synthesis, we generated clonal knockout cell lines that lack P4HA3 and cells that have P4HA3 as the only P4H isoform (*P4ha3*^−/−^ and *P4ha1*^−/−^*;P4ha2*^−/−^ cell lines, respectively). First, we used ddPCR to analyze the absolute and relative expression of the P4HA isoforms in the MEFs and MC3T3 cells, (Fig. 4A). Both parental cell populations and single cells clones derived from them had similar expression patterns. In the MEFs, *P4ha1* was the major isoform (69–70%), while *P4ha2* expression contributed 7–15% and *P4ha3* 16–24% to the total amount of *P4ha* mRNA. Similarly, in the MC3T3 cells *P4ha1* was the most abundant isoform (54– 57%), and *P4ha2* and *P4ha3* contributed 6–7% and 37–40%, respectively. Thus, in both cell types *P4ha3* contributed significantly to the total *P4ha* transcripts, suggesting these cell lines are suitable for studying the function of P4HA3. Lipofectamine-based plasmid delivery of CRISPR/Cas9 plasmids and FACS were used to generate single cell clones, and the knockout was verified by Sanger sequencing and western blotting. We chose to establish clonal cell lines as we aimed for cells that do not have residual enzyme activity in null mutants. The DNA sequencing results suggested that all generated clones had bi-allelic variants that lead to a frameshift and a premature stop codon (<u>Fig. S5, Table S6</u>). The lack of functional protein was validated by western blot. As expected, P4HA3 was absent from the *P4ha3*^−/−^ cells (Fig. 4B, 4C) and neither P4HA1 nor P4HA2 were observed in the *P4ha1*^−/−^*;P4ha2*^−/−^ cell lines (Fig. 4B). Potential compensatory changes in the remaining P4HA isoforms were also evaluated. The lack of P4HA3 protein did not result in any increase in the protein amount of P4HA1 or P4HA2 either in MEFs (Fig. 4D) or MC3T3 cells (Fig. 4E). Similarly, the P4HA3 protein amount remained at original level in the *P4ha1*^−/−^ or *P4ha1*^−/−^ *;P4ha2*^−/−^ MEF cells (Fig. 4D). Uncropped images are shown in Fig. S6 and Fig. S7.

**Figure 4.**
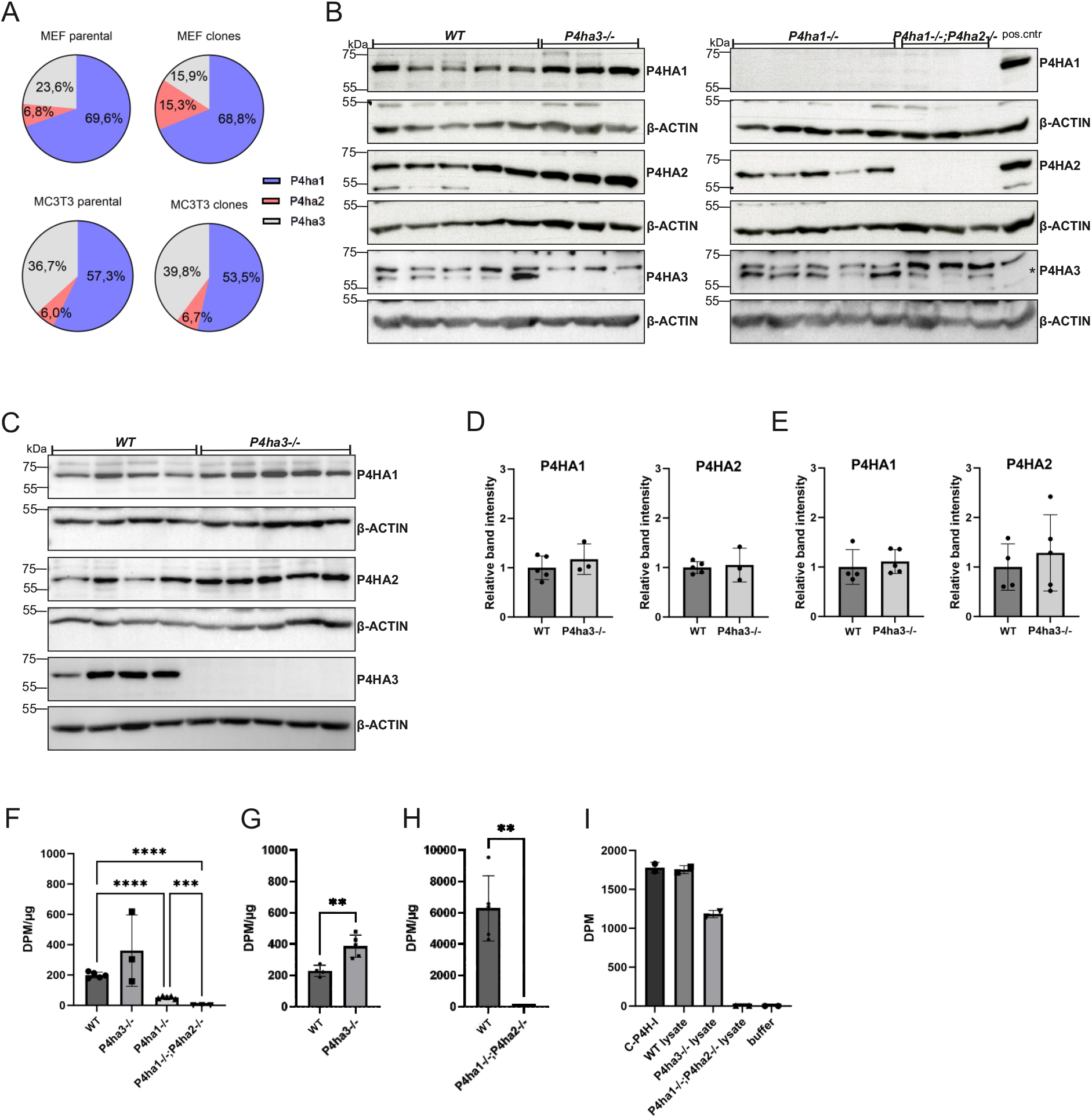
Evaluation of CRISPR/Cas9 edited clonal cell lines. (A) Analysis of the relative expression of the P4ha isoforms in MEF and MC3T3 cells. Droplet digital PCR was used to analyze the relative expression levels of P4ha isoforms in parental and clonal cell lines. The data are shown as mean. For parental samples, 3 technical replicates were analyzed. For the clones, 5 individual clones for MEFs and 4 for MC3T3 cells were analyzed. P4HA knockout verification by western blot of (B) MEF and (C) MC3T3 knockout cell clones. Quantification of P4HA protein amount from (D) MEF and (E) MC3T3 cell clones. The antibodies used in panels B and C are indicated in the right-hand side of the gel images. C-P4H activity was measured in WT and C-P4H deficient (F) MEFs and (G) MC3T3 cells using radiolabeled chick collagen I chains as a substrate. The experiment was repeated with a 12x longer reaction time (6 h) and with more enzyme source material for the (H) *P4ha1*^−/−^*;P4ha2*^−/−^ MEF cell lysates. (I) C-P4H activity was measured in WT and C-P4H deficient MEFs using in vitro transcription/translation produced radiolabeled human α(I) collagen as a substrate. The data are shown in F-H as mean ± SD, n=3–5 individual cell clones per genotype with four technical replicates, and in I data from two technical replicates of the same clones are shown. One-way ANOVA was used when comparing more than three groups and two-tailed student’s t-test when comparing two groups. **P<0.01, ***P<0.001 and ****P<0.0001.

### P4HA3 does not hydroxylate type I collagen polypeptides in vitro

To measure the total C-P4H activity in the generated C-P4H deficient cells, Triton X-100 soluble fractions were used as a source of the enzyme and [^14^C]proline-labelled chick type I procollagen chains as a substrate. The total C-P4H activity was reduced by about 80% in the *P4ha1*^−/−^ MEFs (Fig. 4F). In contrast, neither *P4ha3*^−/−^ MEFs (Fig. 4F) nor *P4ha3*^−/−^ MC3T3 cells (Fig. 4G) did not have any reduction in the C-P4H activity. Interestingly, no detectable C-P4H activity was observed in the *P4ha1*^−/−^*;P4ha2*^−/−^ MEFs (Fig. 4F). The activity assay was repeated with a 12x longer incubation time (6 hours) and with more cell lysate as a source of the enzyme, but still no detectable C-P4H activity in the *P4ha1*^−/−^*;P4ha2*^−/−^ MEFs (Fig. 4H). As the presence of P4HA3 in the *P4ha1*^−/−^*;P4ha2*^−/−^ cell lysate was verified by western blotting using a knockout validated P4HA3 antibody (Fig. 4B), the results suggest that type I collagen is not hydroxylated by the remaining isoform P4HA3 in the *P4ha1*^−/−^ *;P4ha2*^−/−^ cells. We also produced the human type I collagen α(I) chain using *in vitro* transcription/translation, but it also did not function as a substrate for P4HA3 in the lysate (Fig. 4I).

### C-P4H deficiency does not affect the expression of collagens and other collagen synthesis related genes

Next, we evaluated if C-P4H deficiency affects the mRNA expression levels of collagens or other genes related to collagen biosynthesis. The results indicated that loss of C-P4Hs did not affect the mRNA levels of the remaining *P4ha* genes (Fig. 5A, 5B). In the C-P4H knockout cells, the mutated *P4ha1 and P4ha2* transcripts likely undergo a nonsense mediated mRNA decay as the corresponding transcripts were not detected. *P4ha3* transcripts were not completely abolished in the *P4ha3*^−/−^ cells (Fig. 5A, 5B), but western blot analysis showed that there was no P4HA3 protein produced (Fig. 4B, 4C) as expected due to frameshift and early stop-codon (Fig. S5). We also analyzed the mRNA levels of other collagen modifying enzymes, such as *Plod1-3* encoding the three lysyl hydroxylase isoenzymes, *Lox*, and *Lepre1, Leprel1* and *Leprel2* encoding the three P3H isoenzymes (Fig. 5C, 5D). No statistically significant changes were observed in the mRNA levels of these genes. The mRNA levels of a variety of different collagen genes were also evaluated (Fig. 5E, 5F). No statistically significant changes were observed in the mRNA levels of any collagen genes analyzed (*Col1a1, Col1a2, Col3a1, Col4a1, Col4a2, Col4a5* and *Col12a1*). We also observed that the MEFs and MC3T3 cells express very little, if any, *Col4a3, Col4a4, Col9a1, Col14a1* or *Col20a1* genes (data not shown). In addition, the C-P4H deficiency did not affect the mRNA level of fibronectin (*Fn1*) (Fig. 5E, 5F). In conclusion, no alterations in collagen gene expression related to C-P4H loss were observed.

**Figure 5.**
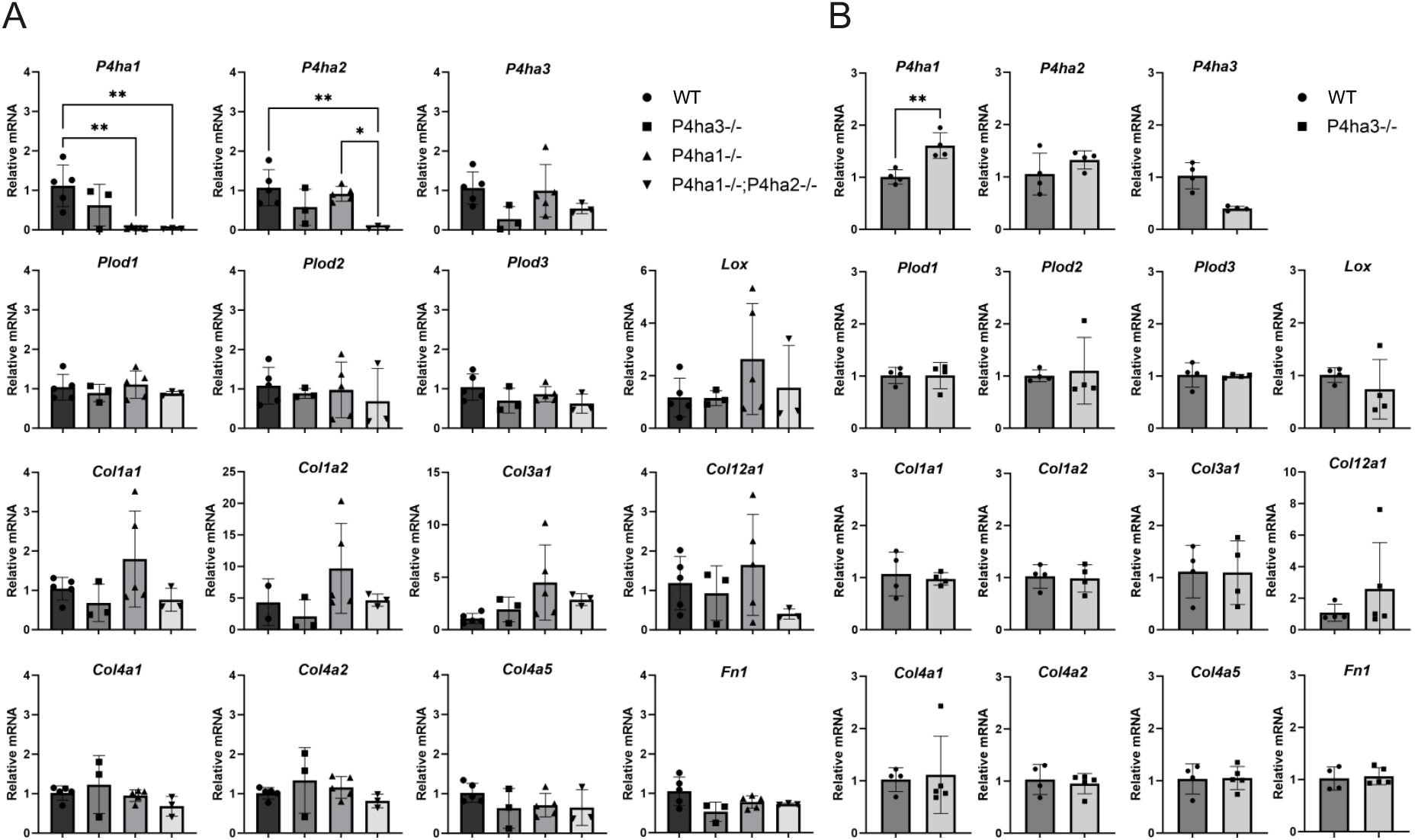
mRNA expression in C-P4H deficient clonal cell lines. Analysis of the mRNA expression of the collagen prolyl 4-hydroxylase isoforms and other collagen related genes in WT and C-P4H deficient A) MEFs and B) MC3T3 cells. The data are shown as mean ± SD, n=3–5 individual cell clones per genotype. Statistical analysis was done with one-way ANOVA. *P<0.05, **P<0.01.

### P4HA3 alone is not sufficient to produce pepsin resistant triple-helical collagen and collagen fibril production

To evaluate how the lack of P4HA3 or the combined lack of P4HA1 and P4HA2 affect collagen synthesis, WT, *P4ha3*^−/−^, *P4ha1*^−/−^ and *P4ha1*^−/−^;*P4ha2*^−/−^ MEFs, and WT and *P4ha3*^−/−^ MCT3T cells were grown in a serum-free medium supplemented with ascorbic acid. The secreted collagen was then extracted from the medium with pepsin digestion followed by salt precipitation. No obvious changes in the amount of secreted pepsin-resistant triple-helical collagen between the WT, *P4ha1*^−/−^ and *P4ha3*^−/−^ MEFs were noticed, whereas no detectable pepsin-resistant collagen was found from the culture medium of the *P4ha1*^−/−^;*P4ha2*^−/−^ MEFs (Fig. 6A, 6C). Similarly, there was no change in the amount of pepsin-resistant secreted collagen between the WT and *P4ha3*^−/−^ MC3T3 cells (Fig. 6B, 6D). Interestingly, some MEF cell clones produced homotrimeric type I collagen molecules (Fig. 6A). The collagen samples from the WT and *P4ha3*^−/−^ MEFs and the MC3T3 cells were further evaluated. CD analysis of collagen showed that the lack of P4HA3 protein had no statistically significant effect on the T_m_ of the collagen triple helix, the T_m_ values being +37.1°C for WT and +37.8°C for *P4ha3*^−/−^ (Fig. 6E). Amino acid analysis was used to measure the amount of 4Hyp and showed that the number of 4Hyp residues in the collagen molecules was not changed (Fig. 6F). This provides further evidence that the lack of P4HA3 does not affect the 4Hyp level in type I collagen. This conclusion was further supported by analyzing the collagen using MS. The data indicated that while the lack of P4HA1 led to a robust reduction in 4Hyp, the absence of P4HA3 did not have any detectable effect (Fig. 6G).

**Figure 6.**
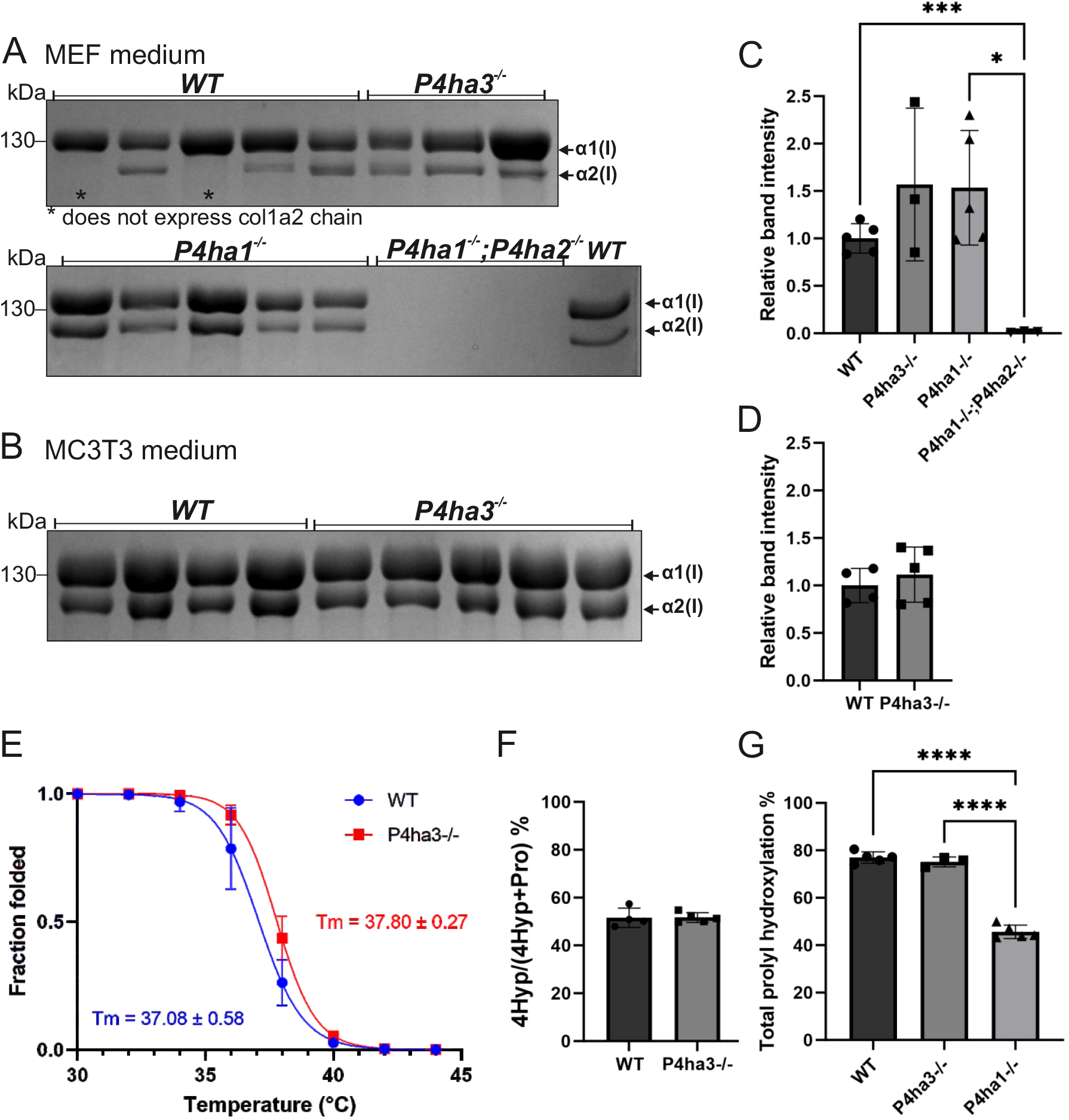
P4HA3 is not involved in type I collagen synthesis. WT and C-P4H deficient MEFs and MC3T3 cells were grown in serum-free medium with ascorbate. Type I collagen was extracted from the culture medium of A) MEFs and B) MC3T3 cells by pepsin digestion followed by salt precipitation. Extracted collagen samples were analyzed by 6% SDS-PAGE under reducing conditions followed by Coomassie Blue staining. Arrows show the positions of the α1(I) (130 kDa) and α2(I) (120 kDa) collagen chains. Quantification of the collagen bands of C) MEFs and D) MC3T3 cells. E) Circular dichroism analysis of collagen from MC3T3 cell medium. F) amino acid analysis of collagen from MC3T3 cell medium and G) mass spectrometry analysis of total prolyl hydroxylation of the -X-Pro-Gly-sequences in the collagen from MEF cell medium. The data are shown as mean ± SD, n=3–5 individual cell clones per genotype. Statistical analysis was done with one-way ANOVA for MEFs and two-tailed student’s t-test for MC3T3 cells. * P<0.0001.

TEM analysis was then used to evaluate how the C-P4H deficiency affects the collagen fibril assembly (Fig. 7). Cells were cultured in the presence of ascorbate for one week. We then investigated the morphology of the collagen fibrils in the extracellular space and measured their thickness. In MEFs, the average fibril diameter was slightly but significantly smaller in the *P4ha3*^−/−^ samples compared to the WT samples, while there was a marked reduction in the average fibril diameter and a clear shift towards thinner fibrils in the *P4ha1*^−/−^ samples (Fig. 7A-C). Also, clearly fewer fibrils were observed in the *P4ha1*^−/−^ MEF cultures. In the MC3T3 cell cultures, the average fibril diameter was significantly smaller in the *P4ha3*^−/−^ cells compared to the WT cells and there was a shift towards thinner fibrils (Fig. 7E-G). Mean fibril diameters for the WT, *P4ha3*^−/−^ and *P4ha1*^−/−^ MEF cultures were 30±6 nm, 29±7 nm and 24±6 nm, respectively, and for the WT and *P4ha3*^−/−^ MC3T3 cultures 49±7 nm and 42±7 nm, respectively (Fig. 7C and 7G). Interestingly, we could not find even a single collagen fibril in the *P4ha1*^−/−^;*P4ha2*^−/−^ MEF cultures, and only some amorphous material and thin filaments of unknown composition was detected (Fig. 7D).

**Figure 7.**
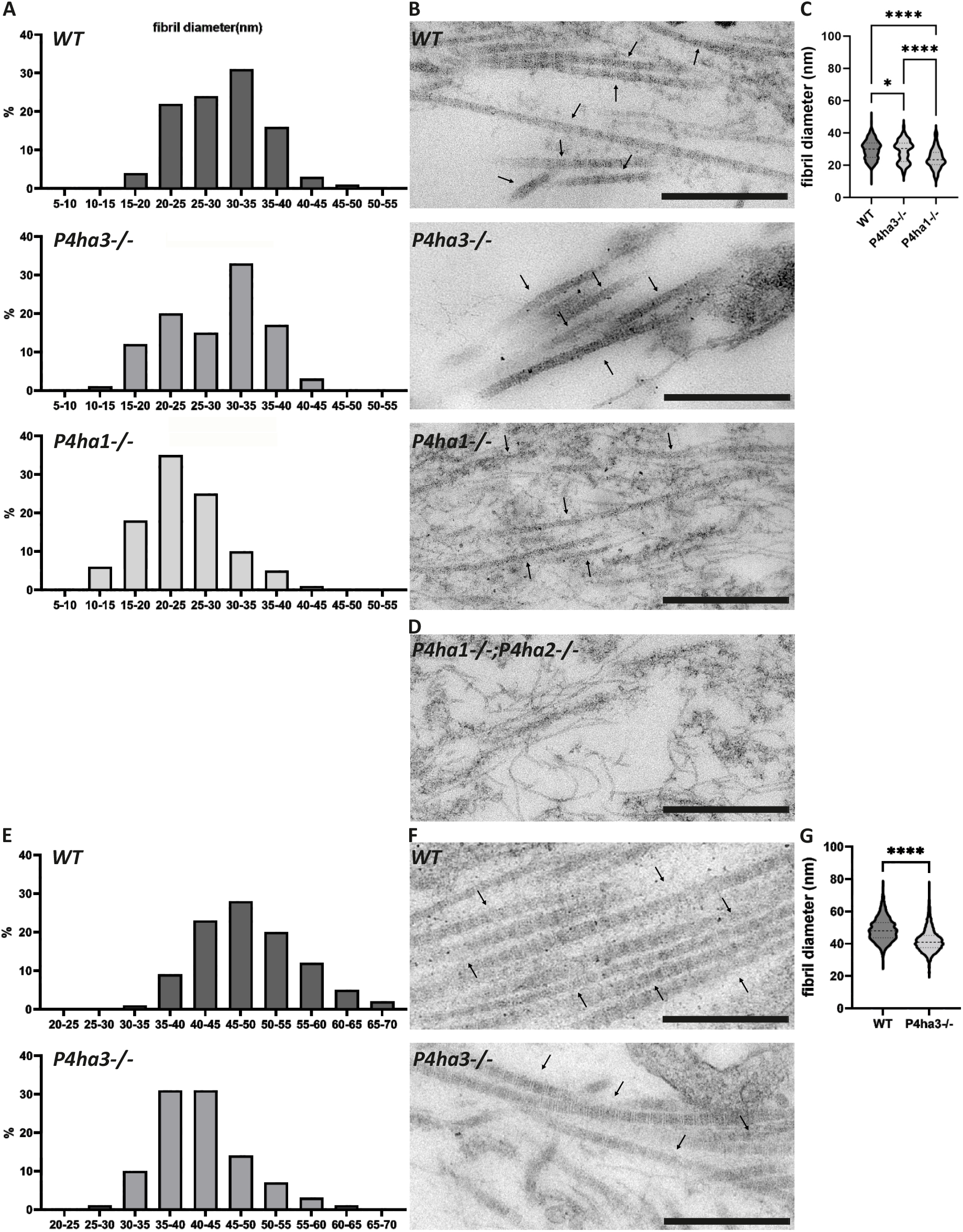
Transmission electron microscopy analysis of collagen fibrils in WT and C-P4H deficient cell cultures. Analysis of the distribution of the collagen fibril diameter as percentage in A) MEF and E) MC3T3 cell cultures. Representative TEM images of B) and D) MEF and F) MC3T3 cell cultures. Mean collagen fibril diameter in C) MEF and G) MC3T3 cell cultures. The data are shown as mean ± SD, the number of analyzed fibrils 92 in MEFs: WT=953 (15 frames), *P4ha3*^−/−^*=*388 (10 frames), *P4ha1*^−/−^=102 (17 frames) and in MC3T3 cells WT=1454 (16 frames) and *P4ha3*^−/−^=1308 (14 frames). Statistical analysis was done with one-way ANOVA for MEFs and two-tailed student’s t-test for MC3T3 cells, *P<0.05, *** P<0.001 and ****P<0.0001. Scale bar is 500 nm in all images.

## Discussion

C-P4Hs have an important role in the biosynthesis of collagens as the formed 4Hyp residues are required for the formation of a native thermostable triple-helical conformation of collagen molecules. The roles of the C-P4H α-subunit isoforms P4HA1 and P4HA2 in collagen synthesis have been widely studied and established (Aro et al., 2015; Holster et al., 2007; Salo et al., 2024; Tolonen et al., 2022), but currently not much is known about the third isoform P4HA3.

Here, we performed the first comprehensive quantitative study of *P4ha* isoform mRNA expression during embryonic and postnatal murine development using ddPCR. *P4ha1* was found to be the major isoform during embryonic development and its expression peaked at 12.5 dpc. Expression of *P4ha3* was also highest at 12.5 dpc, where its expression level was quite similar to that of *P4ha2*. In postnatal tissues, the abundance of all three isoforms was generally highest during the early time points and then decreased by age. Similarly to embryos, the most abundant isoform in postnatal tissues was *P4ha1. P4ha2* expression was lower than *P4ha1* expression, but was detected in virtually all tissues, although the expression was very low in the tail tendon, spleen and liver, for example. The low expression of *P4ha2* expression in tendon has been reported earlier (Wilhelm et al., 2023). In post-natal tissue, the expression level of *P4ha3* was highest from P0 to P4 and then decreased by the two-week time point. The expression level of *P4ha3* was typically similar with, or lower than, *P4ha2*, but in some tissues at certain time points it exceeded *P4ha2* and even *P4ha1* expression. Similarly to our previous study (Tolonen et al., 2022) the highest absolute *P4ha3* expression levels were found in calvaria, (*P4ha3* being the most abundant *P4ha* transcript at P0 and higher than *P4ha2* also at P2 and P4), the calf muscle (higher than *P4ha2* at P0) and the lung (higher than *P4ha2* at P2 and P4 and the most abundant *P4ha* isoform at 1 week of age). *P4ha3* was expressed very little or not at all in 6-week tissues, whereas *P4ha1* was the most abundant isoform at this time point. The results from ISH and scRNA-seq confirmed the widespread expression of *P4ha3* in the developing embryo, including the particularly high expression in the developing skeletal system as also earlier suggested by Tolonen et al. in 2022.

The current data agree with previous studies that have shown that P4HA1 is the major isoform in most tissues and cell types analyzed, whether studied at the mRNA or protein level (Annunen et al., 1998; Kukkola et al., 2003; Nissi et al., 2001; Salo et al., 2024; Wilhelm et al., 2023). Previous semi-quantitative PCR analyses already suggested that the P4HA3 transcript is expressed in many tissues, but in markedly lower levels than those of P4HA1 and P4HA2 (Kukkola et al., 2003). In addition, it has been recently shown by analyzing single cell transcriptomic databases that the P4HA3 transcript is quite rare at a single-cell level and its presence is distinct from P4HA1 and P4HA2 transcripts, which are co-expressed more often (Salo et al., 2024). The data showed that cells expressing the P4HA3 transcript alone or in combination with the other two isoforms were present in very low numbers and were completely absent from the mouse endothelial cell category (Salo et al., 2024). The vast majority of the P4HA expressing cell populations was formed by cells expressing the P4HA1 and P4HA2 transcripts alone or in combination, typically in this order (Salo et al., 2024).

To study the role of P4HA3 in collagen synthesis *in cellulo*, we generated clonal mouse cell lines where the P4HA3 protein is not produced (*P4ha3*^−/−^) and cells where P4HA3 is the only C-P4H isoform (*P4ha1*^−/−^*;P4ha2*^−/−^). We used two different cell lines, MEFs and MC3T3 preosteoblasts, known to produce ample amounts of the main fibrillar collagen, type I collagen, to generate the *P4ha3*^−/−^ cells. These cell lines presented a similar *P4ha* expression pattern with a robust *P4ha*3 expression. *P4ha1*^−/−^ MEFs isolated from the knockout mouse embryos were used to generate *P4ha1*^−/−^*;P4ha2*^−/−^ cells. Inactivation of the target genes was verified by DNA sequencing and immunoblotting. The mRNA levels of the targeted genes were also lower in the knockout cells, suggesting that the introduced frameshift/early stop-codon causing mutations were leading also to the nonsense-mediated mRNA decay. Generally, the lack of P4HA3 did not affect the protein levels of P4HA1 and P4HA2, demonstrating that there was no compensatory reaction. Likewise, the P4HA3 protein amount was not changed in *P4ha1*^−/−^ *or P4ha1*^−/−^*;P4ha2*^−/−^ MEFs relative to the WT cells. Overall, the C-P4H deficiency did not affect the mRNA levels of collagen genes or genes relevant to collagen synthesis. This further supports the previous findings (Salo et al., 2024), that the collagen mRNA expression is not co-regulated with *P4ha* expression.

To analyze the hydroxylation potential and capacity, the amount of total C-P4H activity was measured. Total C-P4H activity was not decreased in the *P4ha3*^−/−^ cells (neither MEFs nor MC3T3 cells). The *P4ha1*^−/−^ MEFs used as a control had less than 30% of the total activity left when compared to the WT cells, which is similar with results reported in previous studies (Holster et al., 2007). Interestingly, no detectable C-P4H activity was found in *P4ha1*^−/−^*;P4ha2*^−/−^ MEFs using radiolabeled chick type I protocollagen substrate. Similarly, when tritiated proline labeled IVTT produced α(I) chain of human procollagen I was used as a substrate, no activity was detected compared to WT and *P4ha3*^−/−^ lysates that produced a robust hydroxylation. This suggested that the single remaining isoform P4HA3 is not able to catalyze the formation of 4Hyp in type I collagen.

Next step was to determine whether P4HA3 has any role in collagen biosynthesis and secretion *in cellulo*. Cells were grown in the presence of ascorbate and triple-helical type I collagen was then extracted from the culture medium with pepsin digestion. Interestingly, we could not detect any pepsin-resistant collagen molecules in the culture media of cells where P4HA3 was the only P4HA isoform. In contrast, no changes in the medium collagen was detected upon the loss of P4HA3. Also, the *P4ha1*^−/−^ cells secreted a similar amount of triple-helical collagen when compared to the WT, although the C-P4H activity was less than 30% relative to the WT, indicating that this activity level is sufficient to maintain type I collagen synthesis in these cells. Collagen samples from the WT and *P4ha3*^−/−^ MEFs and MC3T3 cells were then further analyzed by CD spectroscopy, amino acid analysis and MS. In agreement with the total C-P4H activity results, lack of P4HA3 did not affect the T_m_ of secreted type I collagen molecules and the number of 4Hyp residues was not changed. Since the lack of P4HA3 did not have any effect on these parameters, it further supports the findings that P4HA3 does not contribute to, and is not needed, for the hydroxylation of type I collagen. Indeed, P4HA1 and P4HA2 have been recently shown to have both overlapping and distinct sequence specificity towards different -X-Pro-Gly-triplets and are jointly capable of hydroxylating hydroxylation sites present in type I collagen, P4HA2 having a particular responsibility in the hydroxylation of triplets with negatively charged amino acids in the X-position (-Glu-Pro-Gly- and Asp-Pro-Gly-sites) (Salo et al., 2024; Wilhelm et al., 2023).

We then investigated collagen fibril formation in the C-P4H deficient cells by using TEM analysis. Although there was a statistically significant difference in the collagen fibril diameter between the WT and *P4ha3*^−/−^ cells, the difference was not big, and a similar number of fibrils was observed. We calculated almost 1500 fibrils from both MC3T3 WT and *P4ha3*^−/−^ cells, and the difference in the average diameter was only a few nanometers. We do not know if this difference is of any biological significance. As the P4HA3 substrate is not likely type I collagen and as the type I collagen T_m_ and the number of 4Hyp residues were not altered in collagen from *P4ha3*^−/−^ cells, we could only speculate if P4HA3 affects fibril formation via other mechanisms. In contrast, although the *P4ha1*^−/−^ MEFs were able to produce normal amounts of triple-helical collagen molecules, we detected markedly fewer fibrils with a clear decrease in the diameter in the *P4ha1*^−/−^ cultures relative to the WT cells.

It has been reported earlier that the T_m_ of fibril-forming collagens secreted by the *P4ha1*^−/−^ MEFs is decreased by 1–2°C and the 4Hyp level is reduced by 15% when compared to the WT cells (Salo et al., 2024). Intriguingly, intact *P4ha1* null embryos, from which the *P4ha1*^−/−^ MEF lines were isolated, have been reported to have a slightly increased collagen fibril diameter (Holster et al., 2007). On the other hand, fibril-forming collagens isolated from the *P4ha2*^−/−^ mouse skin have about a 1°C decrease in T_m_ and a 12% decrease in 4Hyp amount. In addition, a reduced fibril diameter has been observed in the *P4ha2*^−/−^ skin when compared to the WT mice (Salo et al., 2024). In conclusion, this study further supports the finding that decreased hydroxylation affects collagen fibril diameter, but the exact mechanisms are yet unknown and there may be other contributing factors. As expected from the lack of pepsin-resistant triple-helical collagen molecules in the culture medium of the *P4ha1*^−/−^*;P4ha2*^−/−^ cells, we did not detect any collagen fibrils in these samples, which further supports the hypothesis that P4HA3 is not involved in the synthesis of type I collagen.

Overall, P4HA3 seems to differ from P4HA1 and P4HA2 in many ways. Human P4HA1 and P4HA2 subunits share a higher sequence identity (65%), whereas P4HA3 subunit is only 35% identical to P4HA1 and 37% identical to P4HA2 subunit (Annunen et al., 1997; Kukkola et al., 2003; Myllyharju, 2003b, 2008). P4HA1 and P4HA2 have been suggested to co-exist in a cell, since they have specific XGP sites they hydroxylate and cannot fully compensate for the loss of another (Salo et al., 2024). The data also showed that cells expressing P4HA3 transcript alone or in combination with the other two isoforms were present in very low quantities and were completely absent from the mouse endothelial cell category, whereas cells expressing both P4HA1 and P4HA2 were the most abundant ones. In line with our studies, P4HA3 is expressed only in few in adult mouse tissues, whereas P4HA1 and P4HA2 are expressed more abundantly.

It is well established that P4HA1 and P4HA2 can efficiently hydroxylate type I collagen. Previously P4HA3 has been suggested to be a collagen prolyl 4-hydroxylase (Kukkola et al., 2003; Van Den Diepstraten et al., 2003) but our findings here do not support those findings at least for type I collagen substrate. Further studies are needed to evaluate if P4HA3 substrate is some other collagen type or even completely non-collagenous protein. Recombinant production and purification of C-P4H-III in sufficient amounts will be important for future studies. Here we also show that cells that lack both established collagen prolyl 4-hydroxylases P4HA1 and P4HA2 (*P4ha1*^−/−^*;P4ha2*^−/−^) are viable, they attach and proliferate on plastic in the absence of properly formed collagen. This cellular model will provide also an interesting model for understanding collagen biology in general. In conclusion, we show that P4HA3 differs significantly from P4HA1 and P4HA2 by having different expression pattern and being functionally distinct by not hydroxylating type I collagen.

## Acknowledgements

Anne Kokko, Eeva Lehtimäki, Raija Salmu and Minna Siurua are acknowledged for expert technical assistance. We thank Dr. Hongmin Tu for CD analyses that were done at the Biocenter Oulu proteomics and protein analysis core facility. MS analyses were performed at the Turku Proteomics Facility. EM sample preparation and imaging was done at the Biocenter Oulu EM core facility (part of Finnish Advanced Microscopy Node of Euro-BioImaging Finland). All the facilities are supported by Biocenter Finland. Mouse maintenance and sample collection was carried out with the support of The Oulu Laboratory Animal Centre Research Infrastructure, University of Oulu, Finland.

## Data availability

The mass spectrometry proteomics data have been deposited to the ProteomeXchange Consortium via the PRIDE (Perez-Riverol et al., 2025) partner repository with the dataset identifier PXD065046 and 10.6019/PXD065046.

## Supporting information

This article contains supporting information

