## Supplementary information for "P4HA3 is abundantly expressed during early embryonic development, but does not catalyze the formation of 4-hydroxyproline in type I collagen"

**List of materials**

Table S1.  Sense- and antisense oligos of CRISPR/Cas9.

Table S2. PCR primers for CRISPR knockout screening.

Table S3. Primers used in qPCR analysis.

Table S4. Mass Spectrometry results

Table S5. Absolute amounts of total *P4ha* mRNA in postnatal tissues  as copies/ng of input RNA.

Table S6. Sequencing results.

Fig. S1. Analysis of the relative expression of the P4ha isoforms in postnatal mouse tissues.

Fig. S2. *P4ha3* in situ hybridization by major systems and structures.

Fig. S3. Dynamics of *P4ha3* expression during embryonic development.

Fig. S4. *P4ha3* expression in skeletal, muscular and cartilage progenitors during embryonic development

Fig. S5. Sanger sequencing chromatograms of A) *P4ha3*$^{-/-}$ MEFs, B) *P4ha1*$^{-/-}$;*P4ha2*$^{-/-}$ MEFs, and C) *P4ha3*$^{-/-}$ MC3T3 cells.

Fig. S6. Uncropped Western blot image of MC3T3s.

Fig. S7. Uncropped Western blot membranes of MEFs.

.

**Table S1. Sense- and antisense oligos of CRISPR/Cas9**

| oligo name | Sequence (5´-3´) |
|---|---|
| P4ha2 guide ex-3-sense | CACCGAGATCAGCTGCCGACCCCGA |
| P4ha2 guide ex-3-antisense | AAACTCGGGGTCGGCAGCTGATCTC |
| P4ha3 guide ex-2-sense | CACCGAACGTCTGCAGTCCGACTGG |
| P4ha3 guide ex-2-antisense | AAACCCAGTCGGACTGCAGACGTTC |

**Table S2. PCR primers for CRISPR knockout screening.**

| oligo name | Sequence (5´-3´) |
|---|---|
| P4ha2 ex3 gDNA F | TGAGAGTTCCCAGCTCTTACC |
| P4ha2 ex3 gDNA R | CCAAATCTGGATCCTCGGCA |
| P4ha3 ex2 gDNA F | AGCCAGCTCCTGATTTTAGCA |
| P4ha3 ex2 gDNA R | GGAAGGCCATGATGTTTTGCA |

F: Forward R: Reverse

**Table S3. Primers used in qPCR analysis**

| Gene | Forward sequence (5´--> 3´) | Reverse sequence (5´--> 3´) |
|---|---|---|
| **Tbp** | GAATATAATCCCAAGCGATTTG | CACACCATTTTTCCAGAACTG |
| **Actb** | AGAGGGAAATCGTGCGTGAC | CAATAGTGATGACCTGGCCGT |
| **Gapdh** | TGTGTCCGTCGTGGATCTGA | TTGCTGTTGAAGTCGCAGGAG |
| **P4ha1** | ATCTCTGACGCGGAGATTGAG | CAGGGTCTTCGTAGCCAGACA |
| **P4ha2** | GTCCTGGTTCGGTGTCCTGAGC | CAGATCGGTCATGTGCCCAATGG |
| **P4ha3** | AGGCCCAACGTACCCCACCT | GTGTGTTGGCTGGGAGCCCA |
| **Plod1** | GATGTCTACTGGTTCCCCATCTTC | CCTGGATCCGGTTGTCCTTATTAT |
| **Plod2** | GCCAGTGGCAATTAATGGAA | CCTGTGTCCATGAGTTTGGA |
| **Plod3** | CCTTCACAACAGCGAGGTGTA | CTCTGGCCCCACCAGCTTTA |
| **Lox** | QuantiTect Mm_Lox_1_SG(QT00098028) | |
| **Col1a1** | TGTGTGCGATGACGTGCAAT | GGGTCCCTCGACTCCTACA |
| **Col1a2** | AAGGGTGCTACTGGACTCCC | TTGTTACCGGATTCTCCTTTGG |
| **Col3a1** | CTGTAACATGGAAACTGGGGAAA | CCATAGCTGAACTGAAAACACC |
| **Col4a1** | TCCCGGACTTCCTGGTATT | CCTTTGTACCGTTGCATCCT |
| **Col4a2** | AGAAAGGAGCAAGGGGTCAG | CTTTGCGCCCTGTAGTC |
| **Col4a3** | GGGAGTGGCAAGAAGTATGG | CGAGATCCCTTCTCAGGAAA |
| **Col4a4** | GGGAGTGGCAAGAAGTATGG | CGAGATCCCTTCTCAGGAAA |
| **Col4a5** | ACGTGGATTTCCAGGCAGT | CTGGTTCCCCCTTCATACCT |
| **Col9a1** | ACCGACCAGCACATCAA | AGGGGGACCCTTAATGC |
| **Col12a1** | AAGTTGACCCACCTTCCGAC | GGTCCACTGTTATTCTGTAACCC |
| **Col14a1** | CCCCAGAATAGAGTGGCACTT | CAAAGCAAGACCTGTTAGGGTAT |
| **Col20a1** | GTCCGACACCTGACTTTCTCA | GCTGAAGTAAGGTTCCCAGGA |
| **Fn1** | ATGTGGACCCCTCCTGATAGT | GCCCAGTGATTTCAGCAAAGG |

**Table S4. Mass Spectrometry results**

| Sample | OG or PG | OG not PG | OG not PG / OG or PG |
|---|---|---|---|
| P4ha3-/- | 1618 | 1302 | 0.80 |
| P4ha3-/- | 1590 | 1228 | 0.77 |
| P4ha3-/- | 1510 | 1144 | 0.76 |
| WT | 1404 | 1038 | 0.74 |
| WT | 1737 | 1344 | 0.77 |
| WT | 1329 | 968 | 0.73 |
| WT | 1321 | 1001 | 0.76 |
| WT | 1578 | 1213 | 0.77 |
| P4ha1-/- | 1364 | 590 | 0.43 |
| P4ha1-/- | 1351 | 635 | 0.47 |
| P4ha1-/- | 1338 | 586 | 0.44 |
| P4ha1-/- | 1281 | 640 | 0.50 |
| P4ha1-/- | 1297 | 573 | 0.44 |

**Table S5. Absolute amounts of total *P4ha* mRNA in postnatal tissues as copies/ng of input RNA. Data are presented as the mean value of combined (Bold) and individual *P4ha1, P4ha2* and *P4ha3* amounts, n=3–5 biological replicates per tissue. Green >500, yellow 100–500 and red <100 copies/ng of input RNA.**

Total P4ha expression, copies/ng of input RNA

| | P0 | P2 | P4 | 1 wk | 2 wk | 6 wk |
|---|---|---|---|---|---|---|
| **Tibia[1]** | **303,5** (169,7/76,7/57,1) | **799,8** (490,7/185,0/124,1) | **1155,0** (689,0/287,2/178,8) | **280,6** (194,1/74,0/12,5) | **59,2** (42,7/12,3/3,1) | **58,8** (45,0/11,7/2,2) |
| **Growth plate[1]** | **517,3** (196,4/111,1/209,8) | **676,2** (285,5/153,3/237,4) | **749,8** (375,9/164,3/209,6) | **223,4** (119,3/55,9/48,3) | **75,8** (38,0/27,1/10,8) | **0,2** (0,1/0,1/0,0)[2] |
| **Calvaria** | **498,8** (247,5/140,1/111,2) | **2635,3** (1540/720,7/374,7) | **888,8** (644,9/180,2/63,7) | **786,9** (487,7/208,7/99,5) | **341,7** (207,6/106,1/27,9) | **89,5** (62,6/21,1/5,8) |
| **Femur** | | | | | | **15,0** (11,6/2,5/0,9) |
| **Femur distal epiphysis** | | | | | | **0,2** (0,1/0,1/0,02) |
| **Bone marrow** | | | | | | **2,5** (2,3/0,1/0,01) |
| **Lung** | **641,5** (419,6/115,5/106,4) | **1054,3** (661,0/161,5/231,8) | **599,9** (413,9/65,3/120,7) | **569,5** (235,8/44,8/288,99) | **81,6** (55,4/14,5/11,6) | **519,3** (388,7/118,3/12,3) |
| **Eye** | **432,4** (295,2/69,8/67,5) | **1064,4** (884,7/120,5/59,2) | **969,0** (776,0/130,0/63,0) | **509,7** (406,0/71,3/32,3) | **900,1** (742,7/132,5/24,9) | **32,1** (24,0/7,8/0,3)[3] |
| **Kidney** | **318,0** (238,4/40,2/39,4) | **613,8** (492,7/87,0/34,1) | **452,5** (331,0/88,4/33,1) | **517,6** (429,3/78,5/9,7) | **397,8** (319,5/75,9/2,4) | **268,7** (222,7/45,8/0,2) |
| **Brain** | **194,1** (123,9/33,5/36,8) | **527,9** (428,0/47,7/52,2) | **545,2** (414,3/64,3/66,9) | | | **217,0** (181,5/23,4/12,1) |
| **Heart** | **289,1** (130,4/77,2/81,4) | **731,6** (407,3/182,5/141,7) | **474,3** (278,1/120,3/75,9) | **76,4** (41,5/20,1/14,8) | **121,5** (69,9/50,5/1,1) | **386,7** (288,4/97,2/1,1) |
| **Aorta** | | | | | | **73,6** (53,3/20,2/0,1) |
| **Microvascular endothelial cell** | | | | | | **1168,2** (1024,0/137,8/6,4) |
| **Calf muscle** | **144,8** (76,2/22,8/45,8) | **858,7** (498/189,7/171,0) | **774,3** (422,2/163,6/188,5) | **259,5** (165,7/60,6/33,1) | **169,9** (109,9/55,5/4,6) | **88,6** (76,4/12,1/0,1) |
| **Skin** | **78,1** (44,4/25,3/8,3) | **99,7** (60,6/34,7/4,5) | **9,6** (8,0/1,5/0,1) | **35,6** (13,6/21,4/0,6) | **27,6** (21,1/6,4/0,1) | **48,9** (32,4/14,9/1,6) |
| **Liver** | | **40,7** (32,9/5,1/2,8) | **35,8** (29,5/3,9/2,4) | **14,9** (13,3/1,0/0,6) | **42,6** (37,5/4,2/0,9) | **27,5** (22,5/4,9/0,1) |
| **Spleen** | | **93,4** (81,5/4,1/7,9) | **100,8** (87,7/6,3/6,7) | **8,1** (7,3/0,3/0,5) | **11,3** (9,5/1,3/0,5) | **8,2** (8,0/0,18/0,05) |
| **Intestine** | | **58,6** (45/8,7/4,8) | **8,9** (7,1/1,0/0,9) | **1,1** (0,8/0,2/0,0) | | |
| **Tail tendon** | | | | | **987,7** (940,7,0/15,4/31,6) | **969,9** (960,4 /5,8/3,7) |
| **FDL tendon** | | | | | | **65,1** (45,1/19,9/0,1) |
| **Achilles tendon** | | | | | | **8,6** (5,2/3,0/0,4) |
| **Xiphoid** | | | | | | **65,1** (45,4/17,0/2,6) |

[1]Tolonen et al. 2022, [2]tibia proximal epiphysis, [3]cornea

**Table S6. Sequencing results**

| | | | | Genomic variant | | Protein level consequence | | |
|---|---|---|---|---|---|---|---|---|
| **cell line** | **genotype before editing** | **gDNA target** | **clone** | **allele 1** | **allele 2** | **allele 1** | **allele 2** | **knockout** |
| MEF | WT | P4ha3-exon2 | 4 | 290insC | 290insC | *W98fs* | *W98fs* | P4ha3$^{-/-}$ |
| MEF | WT | P4ha3-exon2 | 7 | 290insC | 290insC | *W98fs* | *W98fs* | P4ha3$^{-/-}$ |
| MEF | WT | P4ha3-exon2 | 10 | 290insC | 290insC | *W98fs* | *W98fs* | P4ha3$^{-/-}$ |
| MEF | P4ha1$^{-/-}$ | P4ha2-exon3 | 1 | 239delG | 230_233delACCC | *G80fs* | *D77fs* | P4ha1$^{-/-}$;P4ha2$^{-/-}$ |
| MEF | P4ha1$^{-/-}$ | P4ha2-exon3 | 6 | 234delC | 233_234delCC | *P78fs* | *P78fs* | P4ha1$^{-/-}$;P4ha2$^{-/-}$ |
| MEF | P4ha1$^{-/-}$ | P4ha2-exon3 | 10 | 234delC | 228_240del | *P78fs* | *A76fs* | P4ha1$^{-/-}$;P4ha2$^{-/-}$ |
| MC3T3 | WT | P4ha3-exon2 | 4 | 292insT | | *W98fs* | | *P4ha3*$^{-/-}$ |
| MC3T3 | WT | P4ha3-exon2 | 5 | 292insT | 291_295delCTGGA | *W98fs* | *D97fs* | *P4ha3*$^{-/-}$ |
| MC3T3 | WT | P4ha3-exon2 | 6 | 291delC | | *D97fs* | | *P4ha3*$^{-/-}$ |
| MC3T3 | WT | P4ha3-exon2 | 8 | 288delC_290A>G | | *S96fs* | | *P4ha3*$^{-/-}$ |
| MC3T3 | WT | P4ha3-exon2 | 9 | 296insG | 293insC_294delGG | *R99fs* | *W98fs* | *P4ha3*$^{-/-}$ |

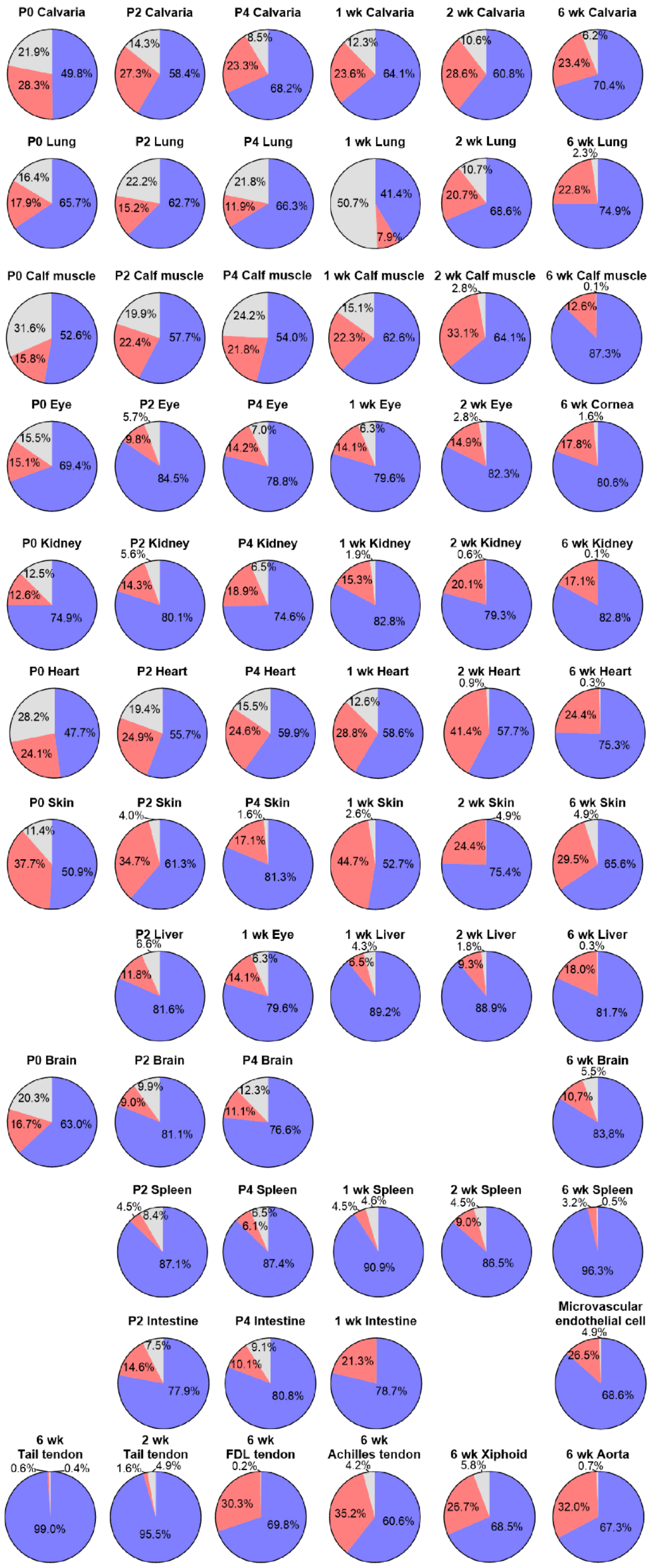


Fig. S1. Analysis of the relative expression of the P4ha isoforms in postnatal mouse tissues. The data used in the analysis are from Fig. 2.

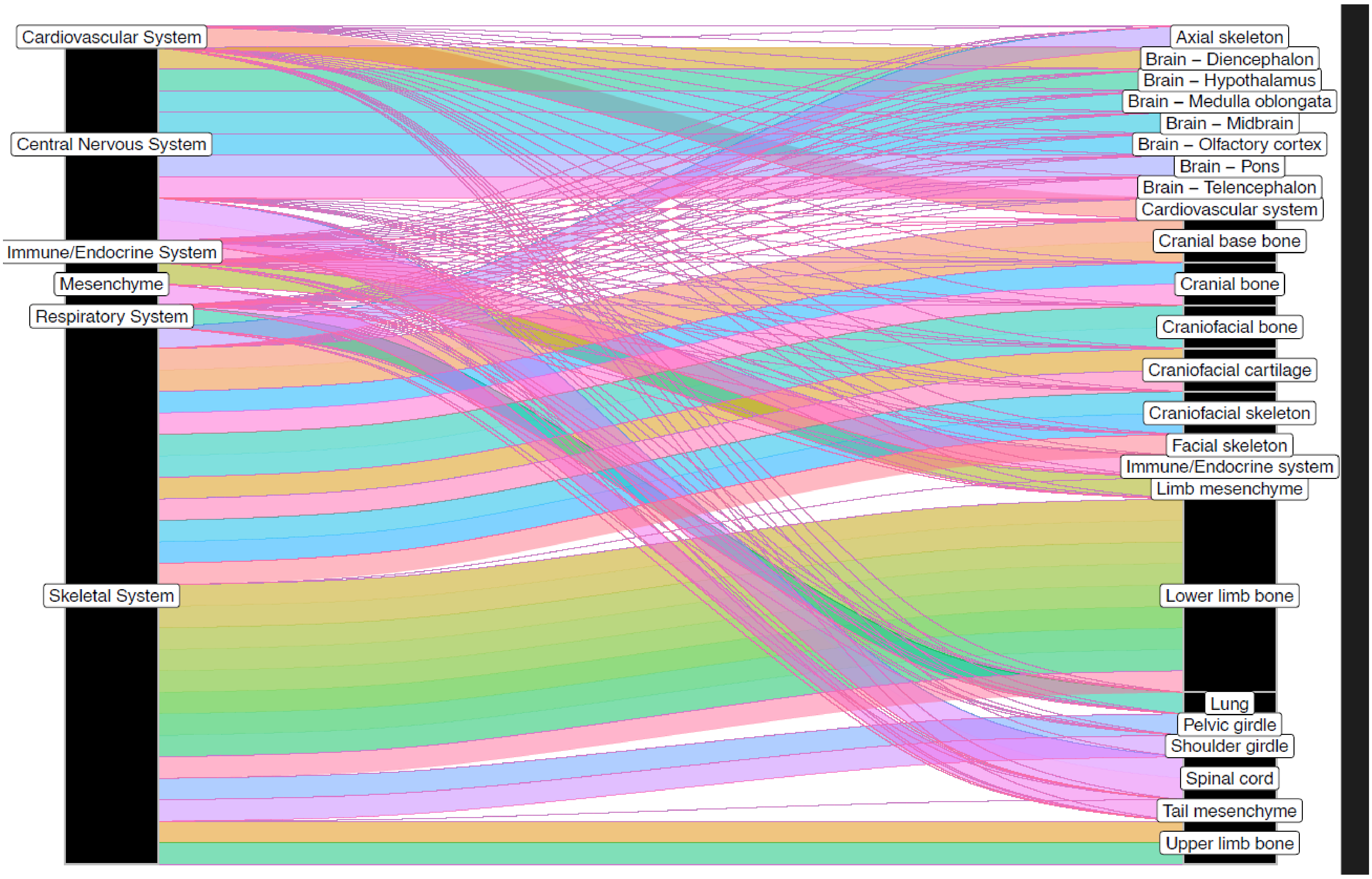


Fig. S2. *P4ha3* in situ hybridization by major systems and structures. Relative quantification of *P4ha3* in situ hybridization (ISH) signal by EMAGE systems and anatomical regions' annotations.

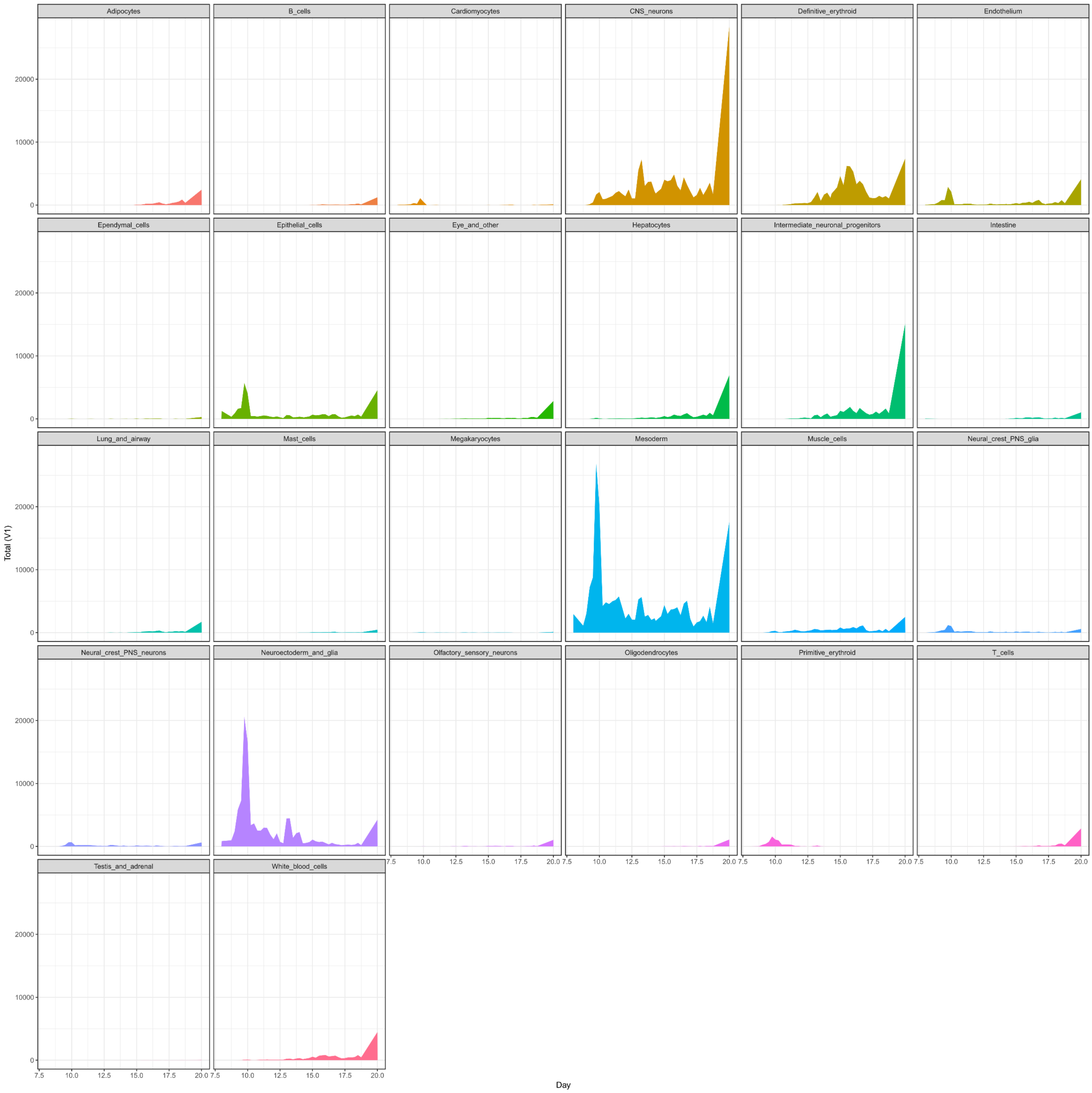


Fig. S3. Dynamics of *P4ha3* expression during embryonic development. Plots show the expression of *P4ha3* (total counts per cell type) across development day, from gastrula to birth.

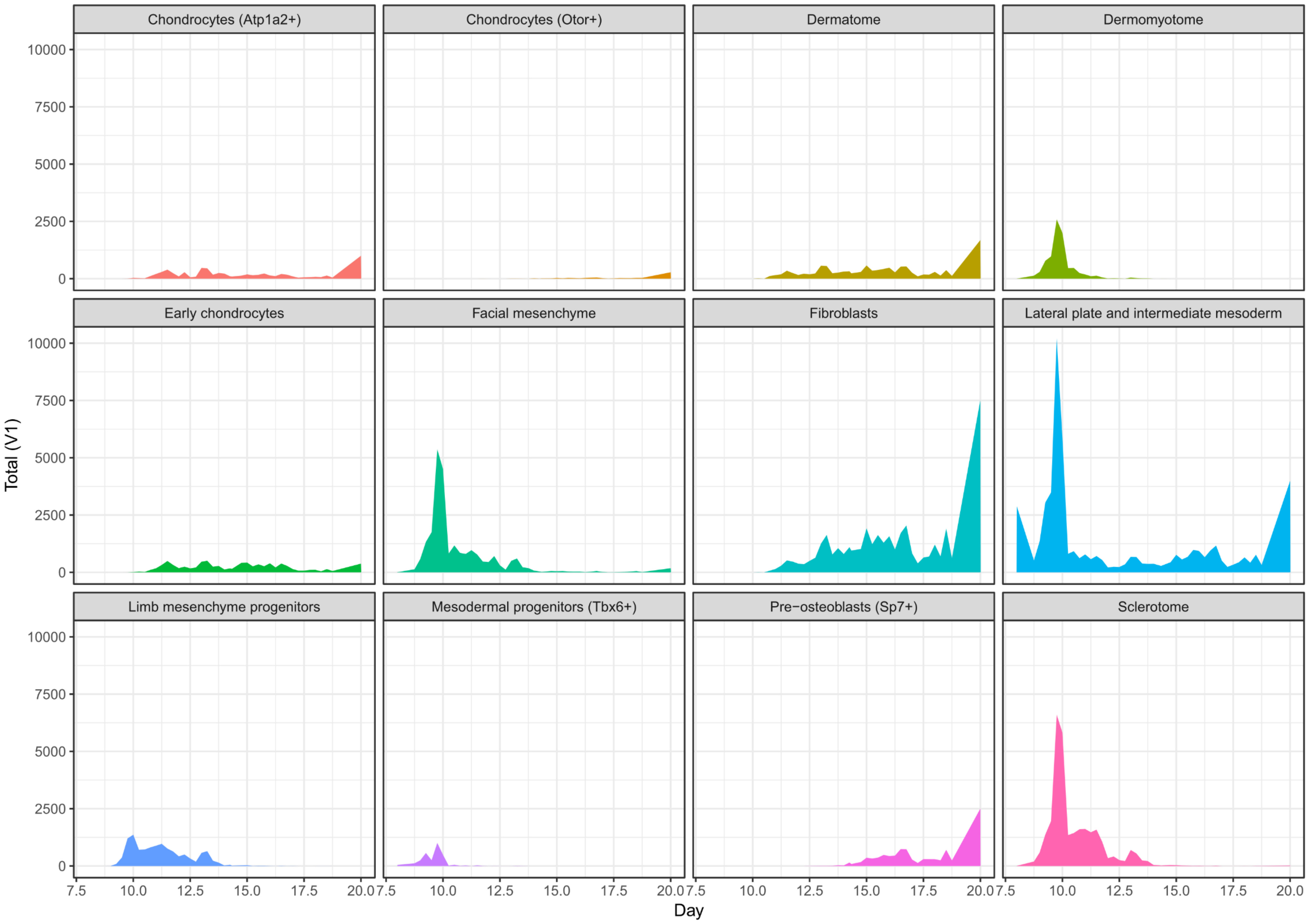


Fig. S4. *P4ha3* expression in skeletal, muscular and cartilage progenitors during embryonic development. Plots show the expression of *P4ha3* (total counts per cell type) across development day, from gastrula to birth.

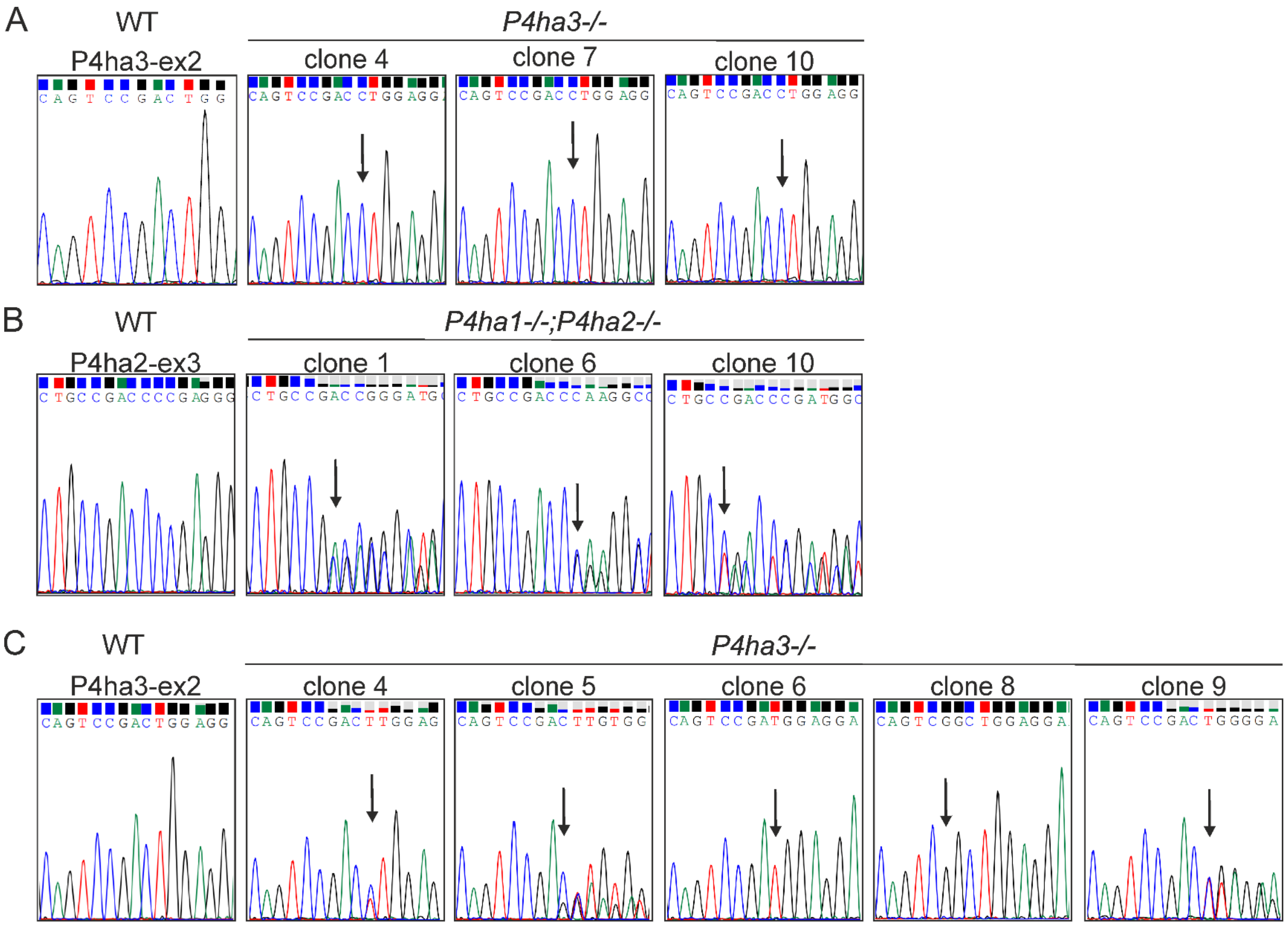


Fig. S5. Sanger sequencing chromatograms of A) *P4ha3*$^{-/-}$ MEFs, B) *P4ha1*$^{-/-}$;*P4ha2*$^{-/-}$ MEFs, and C) *P4ha3*$^{-/-}$ MC3T3 cells. Arrows indicate the targeted site, where the gene edit has occurred. All clones have a frameshift mutation that lead to premature stop codon.

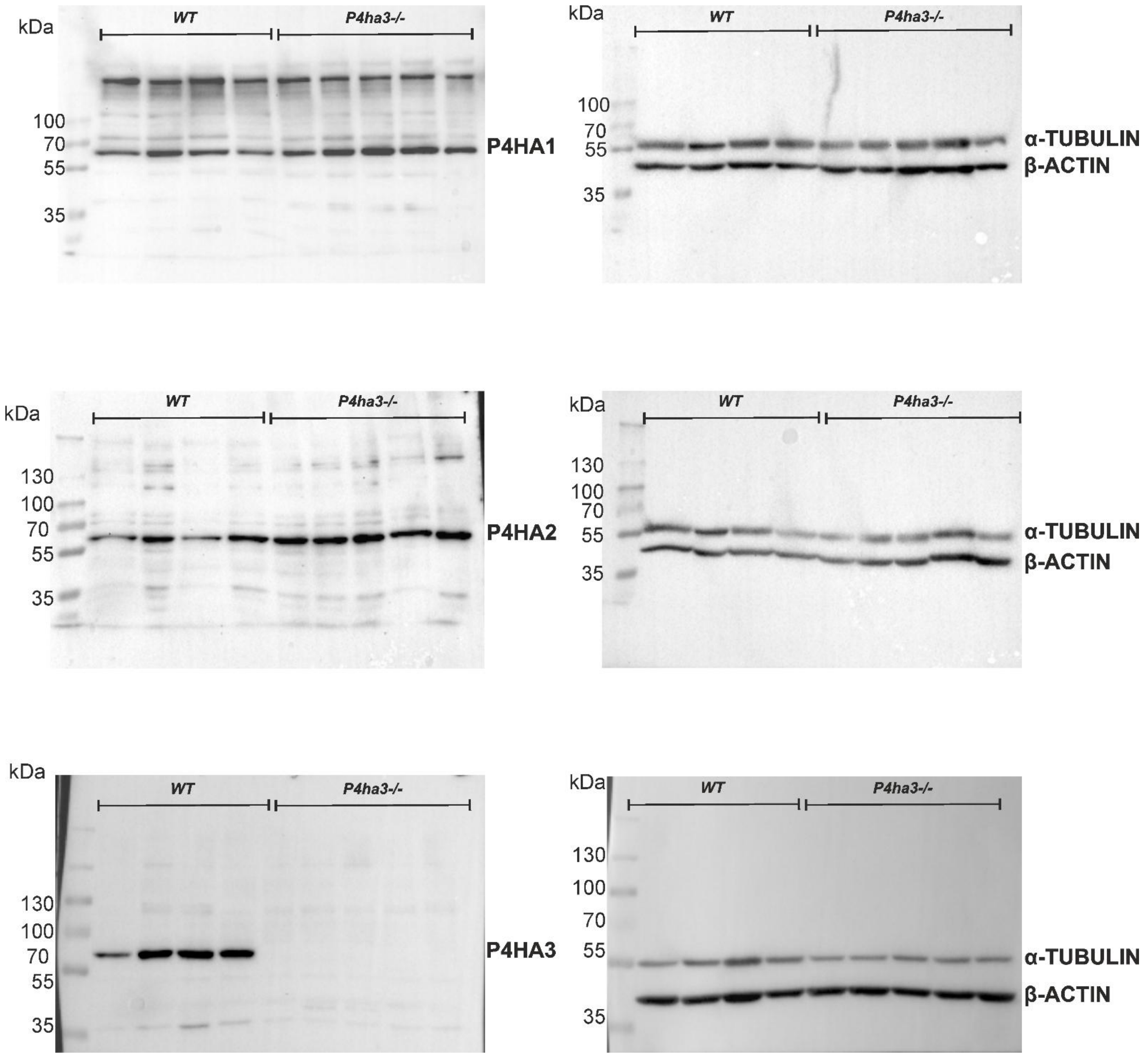


Fig. S6. Uncropped Western blot image of MC3T3s. Whole blot image that was used in Figure 3C.

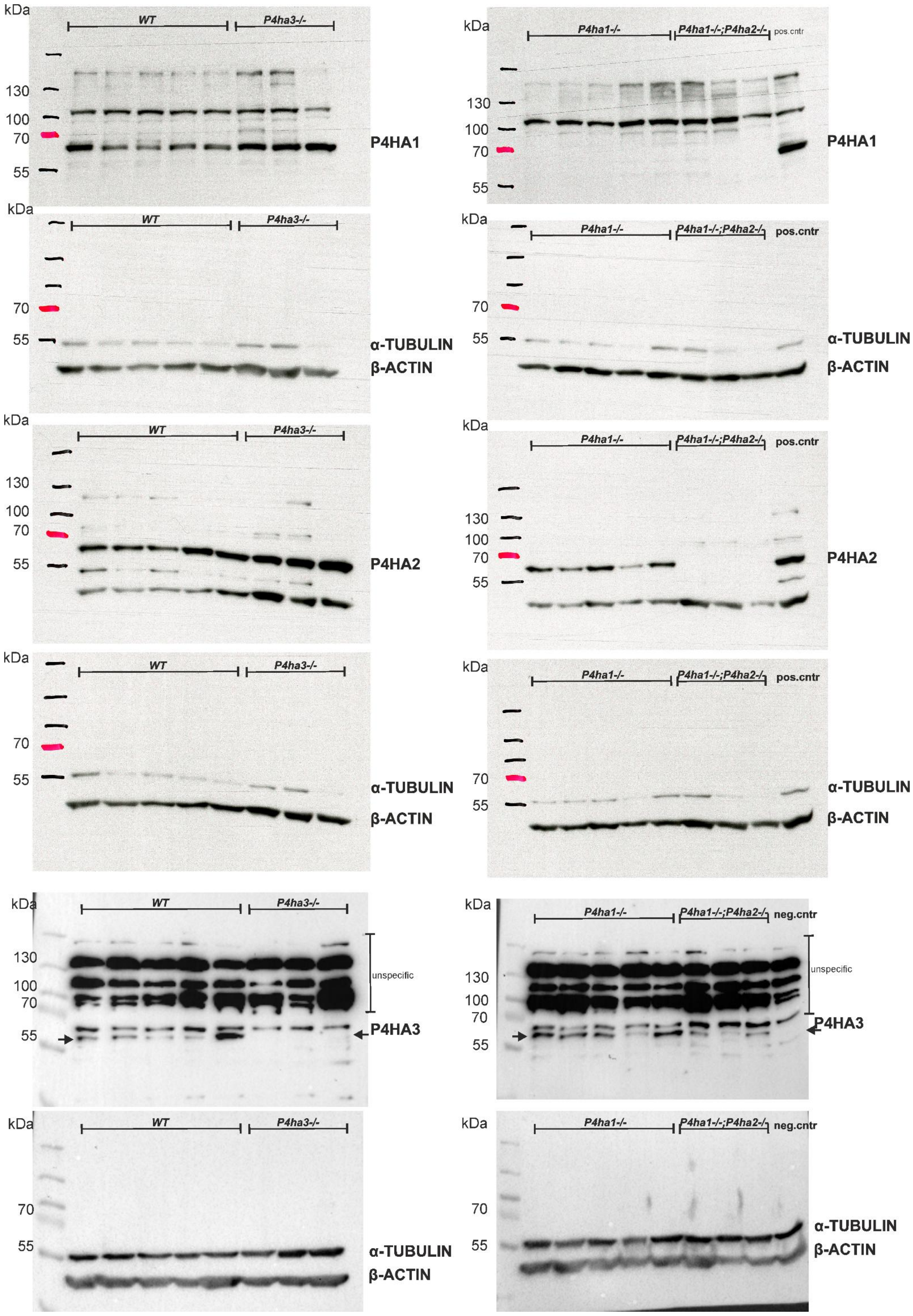


Fig. S7. Uncropped Western blot membranes of MEFs. Whole blot image that was used in Figure 3B.
